# Maintained imbalance of triglycerides, apolipoproteins, energy metabolites and cytokines in long-term COVID-19 syndrome (LTCS) patients

**DOI:** 10.1101/2023.01.13.523998

**Authors:** Georgy Berezhnoy, Rosi Bissinger, Anna Liu, Claire Cannet, Hartmut Schaefer, Katharina Kienzle, Michael Bitzer, Helene Häberle, Siri Göpel, Christoph Trautwein, Yogesh Singh

## Abstract

Deep metabolomic, proteomic and immunologic phenotyping of severe acute respiratory syndrome coronavirus 2 (SARS-CoV-2) patients have matched a wide diversity of clinical symptoms with potential biomarkers for coronavirus disease 2019 (COVID-19). Within here, several studies described the role of metabolites, lipoproteins and inflammation markers during infection and in recovered patients. In fact, after SARS-CoV-2 viral infection almost 20-30% of patients experience persistent symptoms even after 12 weeks of recovery which has been defined as long-term COVID-19 syndrome (LTCS). Emerging evidence revealed that a dysregulated immune system and persisting inflammation could be one of the key drivers of LTCS. However, how these small biomolecules such as metabolites, lipoprotein, cytokines and chemokines altogether govern pathophysiology is largely underexplored. Thus, a clear understanding how these parameters into an integrated fashion could predict the disease course may help to stratify LTCS patients from acute COVID-19 or recovered specimen and would help to elucidate a potential mechanistic role of these biomolecules during the disease course. Here, we report an integrated analysis of blood serum and plasma by in vitro diagnostics research NMR spectroscopy and flow cytometry-based cytokine quantification in a total of 125 individuals (healthy controls (HC; n=73), recovered (n=12), acute (n=7) and LTCS (n=33)). We identified that in LTCS patients lactate and pyruvate were significantly different from either healthy controls or acute COVID-19 patients. Further correlational analysis of cytokines and metabolites indicated that creatine, glutamine, and high-density lipoprotein (HDL) phospholipids were distributed differentially amongst patients or individuals. Of note, triglycerides and several lipoproteins (apolipoproteins Apo-A1 and A2) in LTCS patients demonstrate COVID-19-like alterations compared to HC. Interestingly, LTCS and acute COVID-19 samples were distinguished mostly by their creatinine, phenylalanine, succinate, 3-hydroxybutyrate (3-HB) and glucose concentrations, illustrating an imbalanced energy metabolism. Most of the cytokines and chemokines were present at low levels in LTCS patients compared with HC except IL-18 chemokine, which tended to be higher in LTCS patients and correlated positively with several amino acids (creatine, histidine, leucine, and valine), metabolites (lactate and 3-HB) and lipoproteins. The identification of these persisting plasma metabolites, lipoprotein and inflammation alterations will help to better stratify LTCS patients from other diseases and could help to predict ongoing severity of LTCS patients.

**Graphical abstract:** 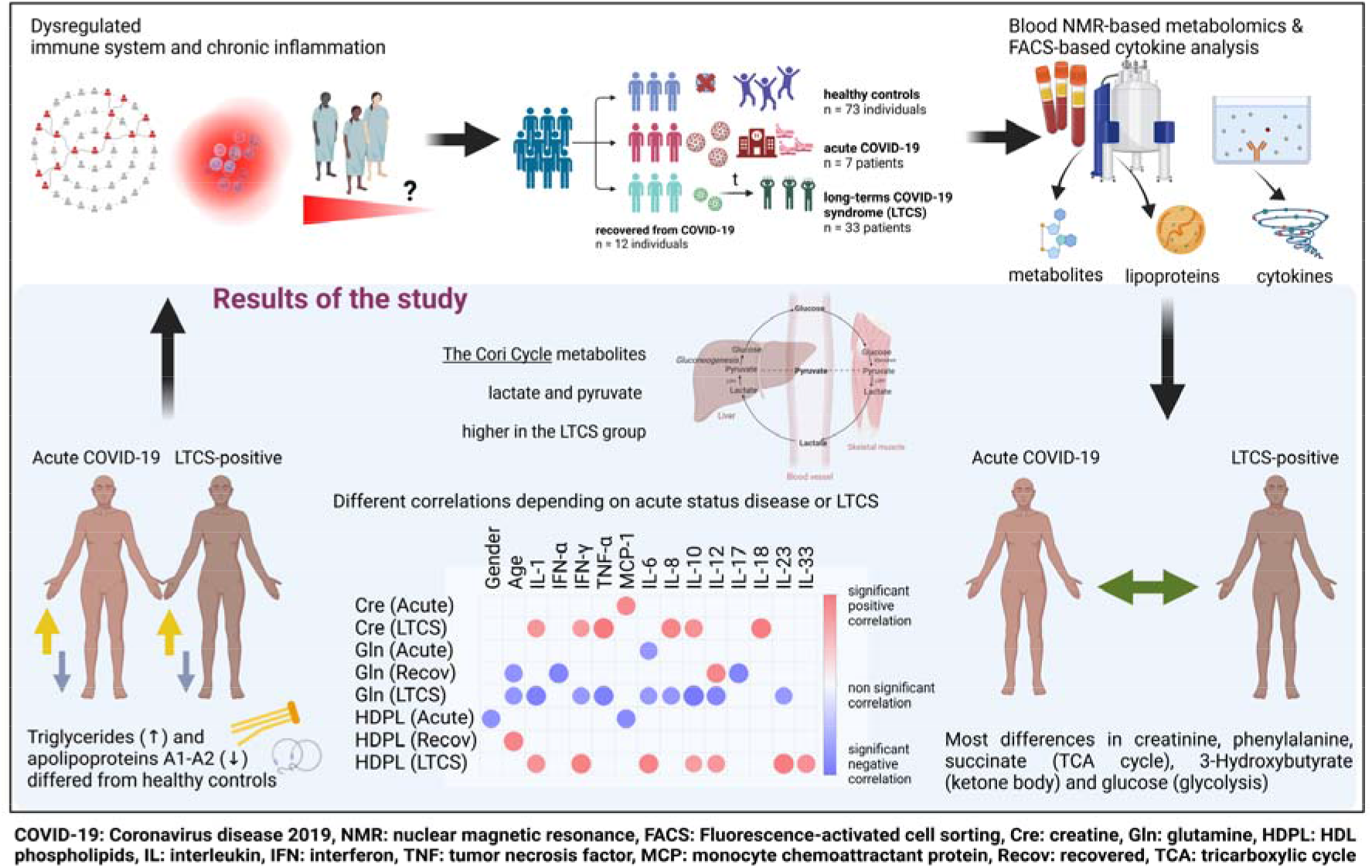

**Layman summary & significance of the research:** Almost 20-30% of individuals infected with the SARS-CoV-2 virus regardless of hospitalization status experience long-term COVID-19 syndrome (LTCS). It is devasting for millions of individuals worldwide and hardly anything is known about why some people experience these symptoms even after 3 to 12 months after the acute phase. In this, we attempted to understand whether dysregulated metabolism and inflammation could be contributing factors to the ongoing symptoms in LTCS patients. Total blood triglycerides and the Cory cycle metabolites (lactate and pyruvate) were significantly higher, lipoproteins (Apo-A1 and A2) were drastically lower in LTCS patients compared to healthy controls. Correlation analysis revealed that either age or gender are positively correlated with several metabolites (citrate, glutamate, 3-hydroxybutyrate, glucose) and lipoproteins (Apo-A1, HDL Apo-A1, LDL triglycerides) in LTCS patients. Several cytokines and chemokines were also positively correlated with metabolites and lipoproteins thus, dysregulation in metabolism and inflammation could be a potential contributory factor for LTCS symptoms.

## Introduction

So far more than 643 million people have been infected with COVID-19 and around more than 6.6 million lives have been lost around the world during the course of the pandemic [1]. Yet, even two years with worldwide SARS-CoV-2 viral infections, the COVID-19 pandemic is still ongoing. Emergence of new variants of concern (VOC) is a great concern despite the development several successful vaccines. Many scientific reports identified the important role of metabolites in the blood serum and plasma of mild, moderate, severe, and recovered COVID-19 patients. In fact, in COVID-19 disease or any other viral infection, immune cells require a lot of energy to fight off the infection, therefore, their metabolic demands drastically increase to produce cytokines and chemokines [2; 3]. A previous study described that peripheral blood mononuclear cells (PBMCs) have a dysregulated glycolysis and oxidative phosphorylation related metabolic profile, with specifically higher lactate and lower glucose levels in mild and moderate COVID-19 patients compared either with healthy controls (HC), convalescent (Co) COVID-19 individuals [4]. Further, specific T cells subsets from acutely infected COVID-19 patients displayed a more extensive mitochondrial metabolic dysfunction, especially cells in CD8 T cell lineages [5]. Further, in vitro activated T cells from acutely infected COVID-19 patients showed a reduced glycolytic capacity and decreased glycolytic reserve, accompanied by relatively low activation of mTOR signalling compared with HC [5]. However, these dysregulated metabolites are released from dysfunctional immune cells as well as tissue damage due to viral infection in the blood [6; 7]. Thus, the detection of metabolites from blood serum or plasma (reservoir and exchanger of metabolites) would give us a hint of the ongoing pathophysiological status of the disease.

Several studies have focussed on how to predict and model the progression of COVID-19 pathological state based on metabolomics and proteomics, including the use of machine learning and mathematical modelling [7; 8; 9; 10; 11; 12; 13]. These studies correlated the metabolites with inflammation and identified that the alterations of several metabolites could be involved in disease progression, with some of them being a direct consequence of the disease. Furthermore, in parallel massive investigative efforts using genomics, transcriptomics, and proteomics to unravel disease mechanisms relevant to SARS-CoV-2 infection were performed on plasma and even fecal samples [14; 15; 16; 17; 18; 19; 20]. A previous study by nuclear magnetic resonance (NMR) spectroscopy identified that lipoprotein subclasses and free cholesterol were increased in both mild and moderate COVID-19 patients, and this study concluded that COVID-19 causes a dysregulation in lipid metabolism, glycolysis, and the tricarboxylic acid cycle [21]. Another NMR study of recovered COVID-19 patients (Recov) after 3–10 months of diagnosis indicated higher plasma cholesterol and phospholipids [22]. Furthermore, lipids changes were determined alongside the metabolites profiling, e.g., amino acids (arginine and glutamine were lower in COVID-19 patients [19]). Additionally, several studies highlighted that inflammatory cytokines such as IL-6 and IL-10 were present in highest levels in severe COVID-19 (acute) compared to moderate/mild or healthy controls (HC) [21; 23; 24].

It is reported that several patients after infection developed a long term COVID-19 syndrome (LTCS) with symptoms such as chronic fatigue, dyspnoea, brain fog, etc. [25]. However, how COVID-19 specific metabolite, lipoprotein and inflammatory mediators relate to the severity of COVID-19 and LTCS outcomes remains poorly understood. Few studies suggested that mitochondrial dysfunction, impaired fatty acid metabolism and cytokine IL-10 production were greatly affected in LTCS patients [22; 26; 27]. Thus, the role of host metabolism and inflammation during the disease progression in LTCS individuals requires further investigation in refined patient cohorts from different geographical regions to validate common and different features of the disease.

Of note, a previously launched in vitro diagnostics research (IVDr) NMR analytical platform demonstrated that for given samples this method can discover quantitative data on metabolite and lipoprotein levels in analysed solutions from either blood serum and plasma (Letertre et al., 2021). The IVDr nMR platform had been already successfully implemented for COVID-19 phenotyping [12; 22; 28; 29; 30; 31; 32; 33; 34; 35]. The samples were collected dated from June 2020 to February 2021 and correspond to the wildtype/alpha mutant of the virus based on epidemiological knowledge. In the current study, we aimed to perform similar investigations on LTCS and control cohorts using ^1^H-NMR based metabolomics, lipoproteome quantification and a targeted multiplex 13-plex inflammation panel. We identified that dynamics of metabolites/inflammation are altered in LTCS individuals.

## Materials and Methods

### Study design and patient recruitments

We used four groups of individuals in this study. The four groups of participants included in this study (Suppl. Table 1) were defined as individuals with: Acute COVID-19 (n=7; with different time points - longitudinal) LTCS (n= 33); Recov (n= 12); and those who lacked any history of positive testing for COVID-19 (n= 73). Recov and LTCS groups were seen in an ambulatory clinical setting. Healthy control samples were recruited for normal blood donation and checked for IgG and IgM antibodies levels to make sure they suffered from no previous SARS-CoV-2 infection (n= 32). Additional healthy control data (n=41) provided by Bruker BioSpin GmbH was generated prior the COVID-19 pandemics. All participants enrolled were of at least 18 years of age. LTCS individuals were patients evaluated at the Tübingen University Hospital for Post-COVID Care between June 2020 and February 2021 for multiomics study cohort (COVID-18 NGS; Ethics number: 286/2020B1 and Clinical Trial number: NCT04364828). They were enrolled only if blood was collected > 28 days after testing positive by SARS-CoV-2 PCR and were experiencing any of symptoms such as fatigue, dyspnoea, brain fog etc. Additional metadata parameters were received and considered for analysis: age and gender status (0 – male, 1 – female, for the purpose of categorization within statistical software). This study was performed in accordance with the Declaration of Helsinki and all patients have been given written consent.

### Sample preparation for the study

Blood samples were collected in the morning at the clinics and delivered to our institute in the afternoon. Initial samples were 9.0 mL EDTA tubes (S-Monovette^®^ K2 EDTA Gel, 9 ml, cap red; Sarstedt, Germany) collected for the isolation of DNA of whole genome and epigenetic study. After completing routine blood tests in the clinical laboratory, the leftover surplus blood from participants who signed informed consents was used for subsequent metabolomics, lipoproteome and inflammation analysis used in this manuscript. The remaining discarded blood samples (2-5 mL) were used for plasma isolation. Plasma separation was performed within 3-4 h after blood collection by centrifuging the blood samples at 2,000 x g for 10 min at room temperature and collected the upper layer. Plasma was stored at – 80°C or until use for both IVDr NMR spectroscopy and 13-plex inflammatory cytokine panel measurements.

### Flow cytometry-based 13-plex inflammatory cytokine assay

To determine cytokine levels from plasma samples obtained from HC, Recov, and acute COVID-19 patients, we employed the LEGENDplex™ Human Inflammation Panel 1 (13-plex) flow cytometry-based assay kit (#740809, BioLegend, San Diego, CA, USA). This panel allowed us for simultaneous quantification of 13 human inflammatory cytokines and chemokines (IL-1β, IFN-α2, IFN-γ, TNF-α, MCP-1 (CCL2), IL-6, IL-8 (CXCL8), IL-10, IL-12p70, IL-17A, IL-18, IL-23, and IL-33). The measurement principle is based on beads which are differentiated from each other based on their size and internal fluorescence intensities on a flow cytometer platform. Each bead set is bound with a specific antibody on its surface and forms capture beads for individual analyte. To detect the cytokine levels, we followed the protocol as recommended by manufacturer’s instruction. Briefly, we first prepared the standard using 1:4 dilution of the top standard (C7) was first prepared as the highest concentration, then serial dilution was done for C6, C5, C4, C3, C2, and C1 by taking 25 μL of diluted standard and added into 75 μL assay buffer. Following, 15 μL of plasma samples were equally diluted with 15 μL assay buffer. Next, 25 μL of the diluted samples were carefully transferred to each well. 25 μL of mixed beads was added to each well. Importantly, beads were mixed well before using by vortex for 30 seconds to avoid bead setting in the bottle. The plate was sealed with a plate sealer, covered entirely with aluminium foil to protect the plate from light, and put on a plate shaker at 800 rpm for 2 h incubation at room temperature. After incubation, the plate was centrifuged at 1.050 rpm for 5 minutes, immediately the supernatant was carefully discarded by flicking the plate in one continuous and forceful motion. The plate then was washed by 200 μL washing buffer. Following 25 μL of detection antibodies were added to each well, the plate was again sealed with a plate sealer, covered entirely with aluminium foil, and incubated for 1 h at room temperature. After incubation, 25 μL of streptavidin-phycoerythrin (SA-PE) was directly added to each well without washing the plate and the plate was sealed and covered in the same manner as described in a previous step. Following the plate was centrifuged for 5 minutes and washed in the same manner as described before. Finally, 150 μL of washing buffer was added to each well and the samples were stored in the cold room until the reading day by flow cytometer.

Data were analysed both manually and automatically by standard curve detection. In automatic gating strategy, two sets of beads were used in this experiment. Each set has a unique size that can be identified by its forward scatter (FSC) and side scatters (SSC) profiles. Based on the internal fluorescence intensities of each set of beads, different resolutions can be achieved. Depending on the type of flow cytometer used, the internal dye was detected via the APC channels. In Beads A there are six bead populations, whereas, in Beads B, there are seven bead populations. The predicted concentration of the cytokine standard levels depicted in different colours. C7 represents the highest level of cytokines, serious dilution taken place among C6, C5, C4, C3, C2, C1 and finally C0 represents the lowest level of cytokines. Log5P analysis were performed to calculate the concentrations of each cytokine for multiple samples based on cloud-based online software provided by BioLegend.

### ^1^H-NMR spectroscopy-based metabolomics and lipoprotein quantification

Raw NMR spectra were recorded using Bruker IVDr (B.I.) methods package for blood samples, which is compatible with EDTA- (ethylenediaminetetraacetate), citrate-, and heparin blood plasma as well as serum samples [36]. The sample preparation was performed following the included standards of procedure (SOP) to ensure reliable results. For quality control, the B.I. BioBank QC™ module was applied. For quantification, the modules B.I. QUANT-PS™ for metabolites and B.I. LISA™ for lipoproteins, respectively, were applied. Blood plasma samples were thawed for approximately 30 minutes at room temperature. An aliquot of 120 μL of each aliquot was pipetted into a 1.5 mL polytetrafluoroethylene (PTFE) container and mixed with 120 μL of commercially prepared pH 7.4 sodium phosphate plasma buffer (Bruker BioSpin GmbH, Ettlingen, Germany). The mixture was then shaken gently for 1 min before transferring 200 μL of it to fill a 3 mm NMR tube (Bruker BioSpin GmbH, Ettlingen, Germany). The autosampler cooling setting was set to 279 Kelvin (4°C). 1D ^1^H-NMR spectra were acquired using a 5 mm triple resonance (TXI; ^1^H, ^13^C, and ^15^N) room temperature probe on a Bruker IVDr Avance III HD 600 MHz system (Bruker BioSpin GmbH, Ettlingen, Germany), which was operated using Bruker’s standard NMR software TopSpin (version 3.6.2). Five one-dimensional ^1^H-NMR spectral experiments were run for each blood sample with water peak suppression and varied pulse sequences to selectively observe molecular components. Firstly, a Nuclear Overhauser Effect SpectroscopY (NOESY) 32-scans NMR experiment was used to show NMR spectrum quality (via the B.I. BioBank QC™) and to enable quantification of metabolites (e.g. glucose, lactic acid, amino acids of the B.I. BioBank Quant-PS™) and high-molecular-weight compounds lipoproteins (as shown in B.I. LISA™). Then, a 32-scan (CPMG Carr-Purcell-Mei-boom-Gill, filtering out macromolecular resonance signals) program was run, as well as 32-scan DIFFusion measurement of, primarily, macromolecular signal massifs (DIFF). Also, a two-dimensional NMR experiment is included within the IVDr methods package and 2-scans J-RESolved spectroscopy (JRES) were recorded to analyse J coupling constants. Additionally, JRES can be useful for a manual data look-up. NMR experiments utilize a group of sample-dependent parameters of frequency offset O1 and duration of 90° pulse P1. Accordingly, regarding the B.I. QUANT-PS™ module, final concentration values as per report pages were used for analysis. The annotation and quantification of serum spectra were provided automatically and server-based by Bruker BioSpin GmbH. Herein, 38 metabolites (via Bruker IVDr Quantification in Plasma/Serum, B.I. Quant-PS™, analysis package) and 112 lipoprotein parameters (via Bruker IVDr Lipoprotein Subclass Analysis, B.I. LISA™, analysis package; Supp. Tab. 2) were identified and quantified in all spectra. As input, final concentrations from B.I. reports were employed.

### Statistical analysis

Statistical analysis was performed with the quantified parameters using the web-based tool MetaboAnalyst 5.0 (Pang et al., 2021). For the software’s analyses, we excluded all features that showed >50% missing values. The remaining missing values were estimated using the feature-wise replacement with 1/5 of a minimum variable value via the Singular Value Decomposition (SVD) computation [37]. The probabilistic quotient normalization (PQN) technique was used to adjust for dilution effects in the corresponding metabolite concentration spreadsheets [38]. To correct for heteroskedasticity, which is not uncommon in this context as concentration magnitudes from metabolites, lipoproteins, and other markers vary strongly, we performed a logarithmic transformation prior statistic. For univariate analysis, volcano plots were generated, which combine p-values generated from unpaired t-tests, and fold change (FC) in one illustration. For figure generation, thresholds for the p-value were established at 0.10 and for the FC at 1.2, respectively. For correlation analyses, we focused on Spearman’s correlation coefficient. Further analyses were conducted using the multivariate approach of unsupervised principal component analysis (PCA) and supervised orthogonal partial least squares discriminant analysis (oPLS-DA). Besides that, PLS-DA was used to assess the discrimination between two groups and identify the parameters that drive this separation. MetaboAnalyst’s biomarker toolbox was used for further biomarker analysis [39]. The univariate analysis (via Mann-Whitney tests), correlational analysis and violin plots, were illustrated using GraphPad PRISM 9.0.1. BioRender.com services were utilized to create some figures within this work.

## Results

### Cohort description and patient demographics

To better interpret the obtained NMR and cytokine data, we first collected the basic metadata from all recruited patients used in the study. We hereby identified in the healthy control group (HC) an age average of 54.4 years, whereas Recov was 67.5 years, LTCS was 56.9 years and acute COVID-19 was 61.1 years (Fig. 1a). Kruskal-Wallis multiple test comparison revealed that the HC group age was significantly less compared with Recov (p=0.022) patients. However, no statistical difference was observed among Recov, LTCS, and acute COVID-19 patients’ age. Gender based analyses were also performed for each group and male and female subjects appeared to be distributed equally in HC and LTCS patients (Fig. 1b & Suppl. Table 1). Further, we identified the LTCS patient sample collection from date of infection to plasma collection for the study. We identified that median of the sample collection was 152 days after viral infection with minimum 47 and maximum 308 days (Fig. 1c), whereas in. the case of Recov patient group it was 128 days (Fig. 1c). LTCS and Recov patients sample collection was significantly different (p=0.03), thus it appeared that LTCS patients and Recov patients had a clear demarcation of the symptoms. Further, we obtained ten major different symptom parameters to define LTCS patients: we identified that our cohort (n=33) had fatigue (>54.5%), dyspnea (>51.5%), dizziness (>21%) as major symptoms (Fig. 1d). Anosmia (>15%), ageusia (>15%), headache (>9%), anxiety (>9%), myalgia (>9%), and neuropathy were less common symptoms (>6%).

**Fig. 1.**
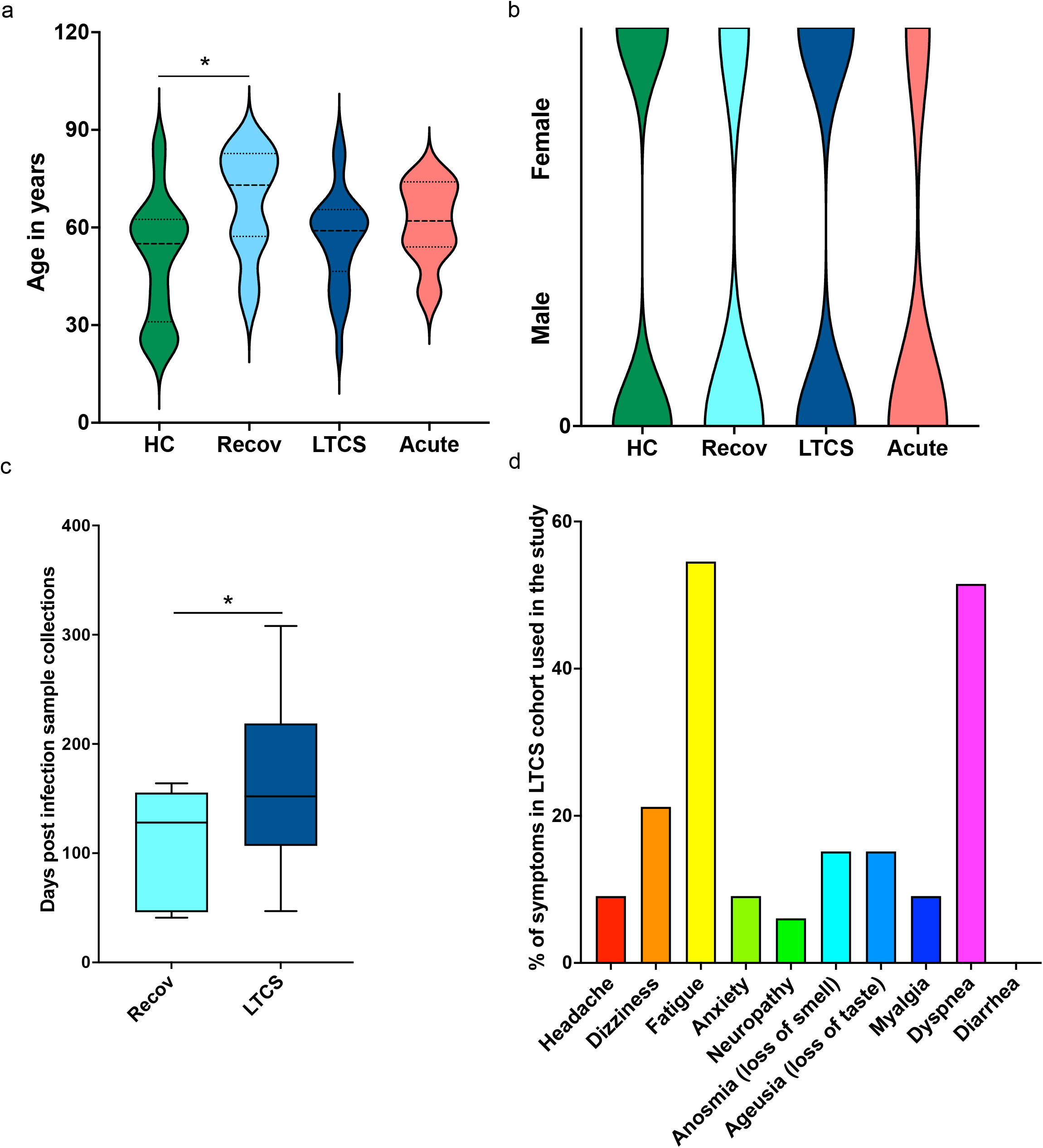
Patient demographics of LTCS patients and study cohort. Age dependency in clinical patient groups and a substantial age difference between healthy controls and recovered patients (sub-plot a). Gender-based structural map for the patient groups (sub-plot b). There is a statistically significant difference between the recovered group and the LTCS group in number of days post infection registered (sub-plot c). The rows graph shows (sub-plot d), in percentages, comorbidities as subclinical cofactors for the LTCS group.

### Dysregulated metabolites in severe and LTCS patients

Several studies identified that blood metabolites are dysregulated especially in severe COVID-19 patients and in recovered patients [3; 39; 40; 41; 42]. This is further affected by different variant strains and collection times [43]. However, information on how metabolites and inflammation affect for LTCS patients has started to emerge only recently [22; 26]. In our study, we used quantitative IVDr ^1^H-NMR spectroscopy to distinguish metabolites levels in HC, Recov, LTCS, and acute COVID-19 patients. First, we confirmed QC check using B.I. QUANT-PS™ analysis method and identified that only 84 out of 89 passed strict quality check and similarly Recov 12/14, LTCS 33/54, and all acute COVID-19 patient was used with different time points (Fig. 2a). We first compared entire cohort of samples with different groups based on quantifiable metabolites data (B.I. QUANT-PS™) and untargeted principal component analysis (PCA). We found that the acute COVID-19 patient group showed a clear separation with either with LTCS, Recov or HC group (Fig. 2b). PCA loadings and the graphical representation of metabolic parameters contributed was performed as well (Suppl. Fig. 1). Further, PLS-DA’s variables in projection importance score plot (VIP) loading rating suggested that the amino acid creatine and a ketone body 3-hydroxibutryrate were present at highest level in the plasma samples of acute COVID-19 patient whilst citrate and histidine were present in LTCS patients among all other groups (Fig. 2c). Recov patients had the highest amount of pyruvate and lactate levels (Fig. 2c-d), as illustrated also on the heat map plot. Further, we identified that formate, acetone, and citrate were present in higher amount in LTCS compared to Recov patients (Suppl. Fig. 2). Due to the limited number of samples in the Recov and acute group we mostly focus in this paper on the comparison between HC and LTCS. A supervised classification model was built using orthogonal projections to latent structures discriminant analysis (oPLS-DA) to distinguish between HC and LTCS patients, using metabolites as variables. We hereby could observe a clear difference and heightened levels of pyruvate, lactate, methionine and alanine in LTCS patient compared with HC (Fig. 2e-g). The regression analysis also highlighted that pyruvate, lactate and methionine were top on the S plot. Furthermore, the metabolite panel volcano analysis results showing trend-like (FC > 1.2, p ≤ 0.10) changes for lactate, pyruvate, and methionine (up) and phenylalanine, glycine, Gln/Glu (glutamine-glutamate ratio), lysine and acetate (down) in LTCS compared with HC (Fig. 2h). Finally, we compared and revealed the overall changes in the metabolites among different groups (Fig. 2i). Despite the low concentrations of some metabolites, we were able to determine the degree to which the dimethylsulfone was low for different LTCS patients at its average concentrations.

**Fig. 2.**
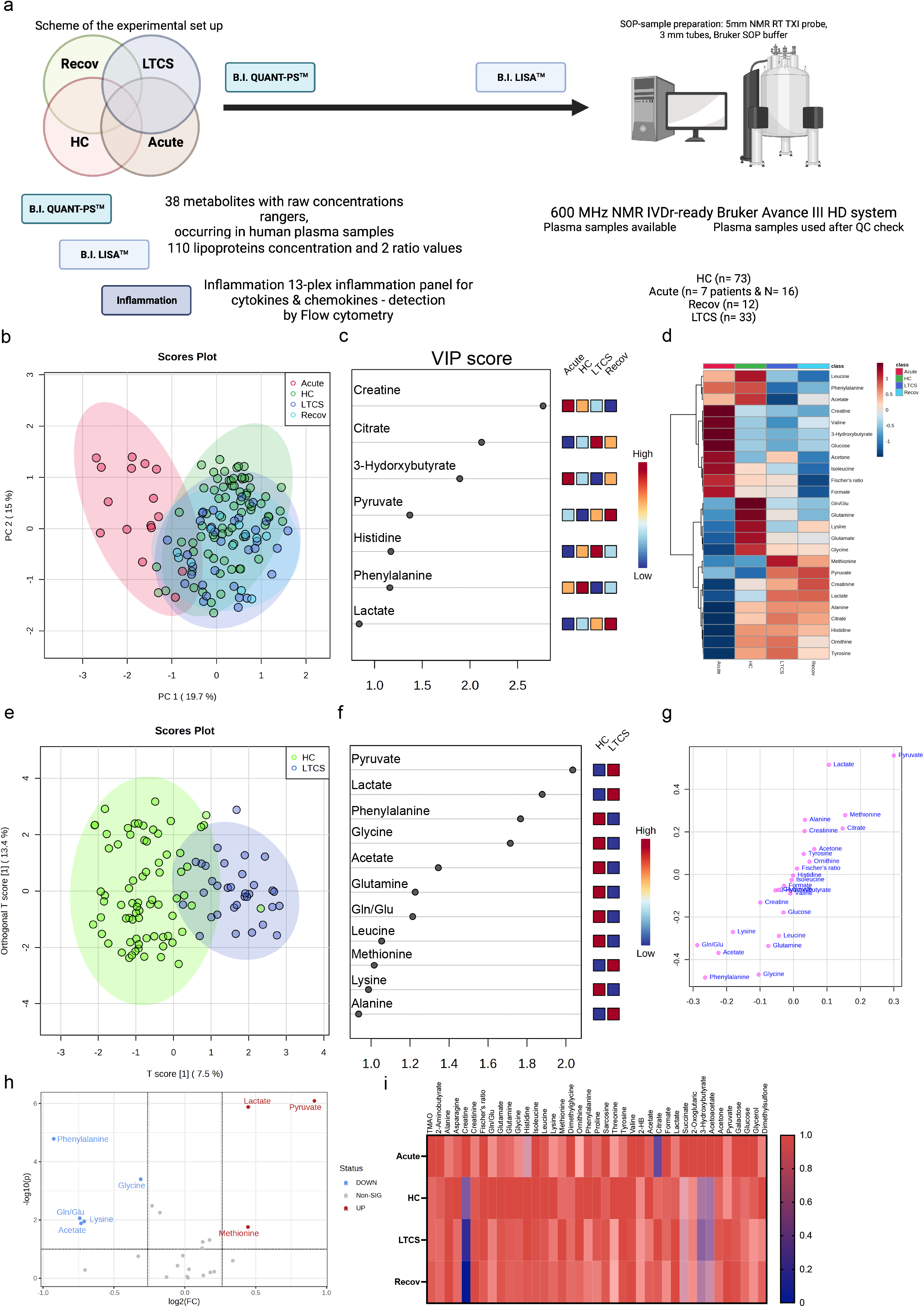
Identification of metabolites in LTCS patients. The image above depicts the study’s methodology (sub-plot a). PCA and PLS-DA studies were performed for the whole cohort data. This analysis was done out based only on quantifiable metabolites data (B.I. QUANT-PS™). The 4-group distribution was shown in (b) using the coordinates of principal components 1 and 2. The values that contributed the most to these VIP scores are shown here by the subplot (c), which are sorted from most significant to least significant. The metabolite panel variables’ average trends were presented by sub-plot (d). oPLS-DA study was performed, and LTCS vs control patients (HC) were compared. This analysis was done out based only on quantifiable metabolites data (B.I. QUANT-PS™). The two-group distribution was shown in (e) using the coordinates of loading components 1 and 2. The values that contributed the most to these VIP scores and S-plot data of the regression model are shown here by the subplots (f) and (g), which are sorted from most significant to least significant. Metabolite panel Volcano analysis results showing trend-like (FC > 1.2, p ≤ 0.10) changes in ratio of LTCS/HC as presented by sub-plot (h). For each patient group (Recov (EDTA plasma) n=12, HC (serum or heparin plasma) n=73, LTCS (EDTA plasma) n=33, Acute (Heparin plasma) n=16 samples), an average normalized (scaled 0 to 1, averages were divided by a maximal average per variable) Heat map analysis conducted by sub-plot (i).

We delineate that LTCS compared with acute COVID-19 patients have a highly significant change in several metabolites including alanine, histidine, citrate, lactate, pyruvate, creatine, succinate, and glucose (Suppl. Table 3). Yet, we have been unable to establish any differences between the LTCS and Recov groups that are statistically significant. This is also not surprising, as the n-number for the Recov group is very small. The examination by a regression model, however, made it possible to zero in on factors that had a hand in the classification of the groups (Suppl. Fig. 2). Elevated levels of formate were detected for the LTCS, but the group also had a tendency of lowered amounts of acetate, creatinine, lysine, valine, pyruvate, phenylalanine, and lactate when compared to the Recov individuals.

Overall, the energy metabolites of citrate and pyruvate were much higher in the LTCS and Recov groups than in the acute COVID-19 patients (Fig. 2 & Suppl. Fig. 3). We next identified pathway alterations. Overall, six pathways were mainly identified which had a significant difference including TCA cycle, ketone bodies, alanine/aspartate/glutamate metabolism, glycolysis, glycine/serine/threonine metabolism, and arginine/proline metabolism. In all six pathways, glycolysis pathway has less abundant metabolites from the acute patient samples. At the same time, TCA cycle metabolites were high in both Recov and LTCS patient groups with high significance levels. Finally, we were able to observe slightly lowered levels of glycolysis metabolites in the LTCS group as well. Thus, our presented data defines that metabolic dysregulation in LTCS and acute COVID-19 spectra of metabolites changes.

### Imbalanced lipoproteins are key characteristics for LTCS

Several studies on mild/moderate and acute COVID-19 patients have implicated the important functions of lipoproteins in disease development [12; 33; 36; 44; 45; 46; 47]. In our study, the four cohort groups were partially separated by the PCA (Fig. 3a, PCA loadings plot – Suppl. Fig. 4). showed that the Recov group was characterized by the highest levels of LDL-5 and LDL-6 subfraction cholesterol content (Fig.3b). Lipoproteins such as V5FC, V5CH, L6TG, and L4PL were increased mostly in LTCS patients (Fig. 3b&c). We also identified that several lipoproteins were present in lower amounts in LTCS compared with Recov patients (Suppl. Fig. 5). Herein, we found that HDL-4 triglycerides were significantly higher in the Recov group compared to the LTCS group. Due to the small n-number of the Recov group we then focused only on HC and LTCS patients. Here we observed in the oPLS-DA plots that HC and LTCS form two clusters though not being entirely separated (Fig. 3d). Variable projection regression analysis revealed that a greater number of lipoproteins were highly abundant in HC compared with LTCS patients (Fig. 3e-f). Based on volcano plots, we identified that 18 lipoproteins were increased whilst 38 lipoproteins were decreased in LTCS patients compared with HC, (Fig 3g). On the Volcano analysis plot, we saw decreases in Apo-A2 and cholesterol parameters (fold changes > 1.2, p values 0.10) and triglycerides LTCS-elevated lipoprotein variables. We detailed the substantial differences in several metabolites between LTCS and severe acute COVID-19 patients, including HDL cholesterol and apolipoprotein B100 Apo-B (Fig. 3h).

**Fig. 3.**
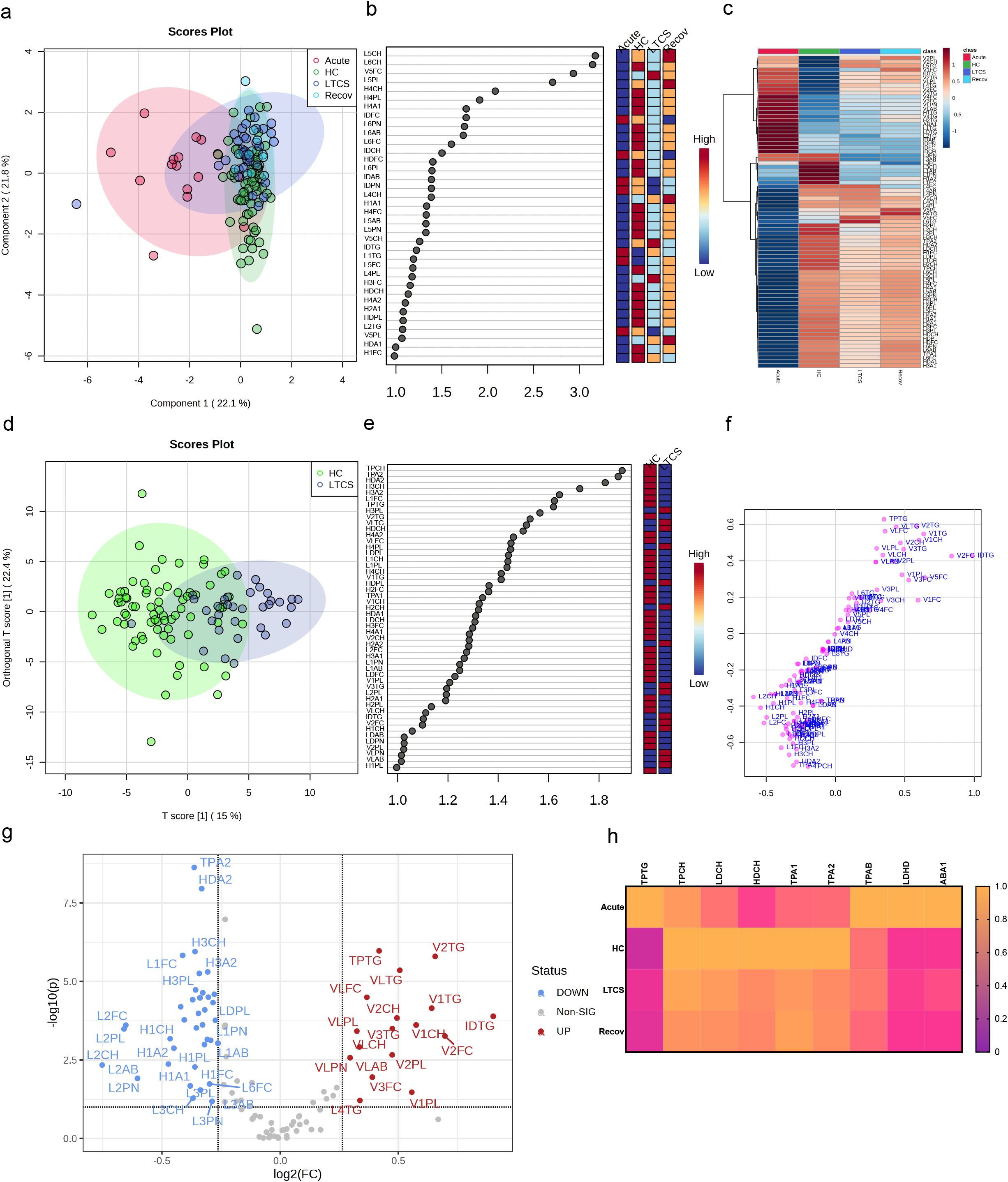
Lipoprotein profiling in LTCS patients. PLS-DA was done out based only on the lipoprotein data panel (B.I. LISA™). The 4-group distribution was shown in (a) using the coordinates of loading components 1 and 2. The values that contributed the most to these VIP scores are shown here by the subplot (b). Lipoprotein data variables’ average trends were presented by sub-plot (c). oPLS-DA study was performed, and LTCS vs control patients (HC) were compared. This analysis was done out based only on the lipoprotein data panel (B.I. LISA™). The two-group distribution was shown in (d) using the coordinates of loading components 1 and 2. The values that contributed the most to these VIP scores and S-plot data of the regression model are shown here by the subplots (e) and (f), which are sorted from most significant to least significant. Lastly, the lipoprotein panel Volcano analysis results showing trend-like (FC > 1.2, p ≤ 0.10) changes in ratio of LTCS/HC as presented by sub-plot (g). Lastly, the main lipoprotein panel variables’ average trends were presented by sub-plot (h). For each patient group (Recov (EDTA plasma) n=12, HC (serum or heparin plasma) n=73, LTCS (EDTA plasma) n=33, Acute (Heparin plasma) n=16 samples), an average normalized (scaled 0 to 1, averages were divided by a maximal average per variable)

Performing an additional analysis based on the Mann-Whitney test (Suppl. Table 4), we identified that Recov (**), acute (****), and LTCS (****) had higher blood triglycerides than the healthy control group, something that has been reported for COVID-positive individuals previously [31; 48; 49]. Moreover, very low-density lipoprotein (VLDL) phospholipids were also higher in Recov (*) and LTCS (****, Suppl. Table 5). Interestingly, free cholesterol levels were not significantly different between LTCS and acute COVID-19 groups. Acute COVID-19 group had the highest blood triglyceride levels versus the Recov (**) and LTCS (***) groups (Suppl. Table 4).

### The combination of metabolites, lipoproteins and cytokines orchestrates pathological phenotypes

Several studies identified that inflammation, metabolism, and lipoprotein content act in unison to overall inform the status of specific disease state such as mild, moderate, or severe in COVID-19 patients and thus can help us to predict and stratify the disease severity [2; 3; 21; 36; 46; 50; 51; 52; 53]. Our cytokines and chemokine profiling showed that acute COVID-19 samples had a trend of highest levels of cytokines & chemokines compared with either HC, LTCS or Recov (Suppl. Table 8). Furthermore, most of the cytokines and chemokines had a tendency of lower levels in either LTCS or Recov compared HC, except IL-18 chemokine which was found to be higher in LTCS and Recov compared with HC, however not reaching a significance level (Suppl. Table 6). We validated previously published data that IL-8 chemokine and IL-6 and IL-10 cytokines were abundantly present in the acute COVID-19 patients [54; 55]. With our data we also performed PCA and PLS-DA analysis and identified a major separation among acute and LTCS patients (Suppl. Fig. 6a). In a correlation analysis of cytokines, chemokines and metabolites revealed that acute COVID-19 patient had the highest levels of cytokines (Suppl. Fig. 6b). The Recov patients showed medium levels compared with LTCS, whereas all these cytokines and chemokines were present in low abundance in LTCS patients in overall comparison (Supp. Fig. 6b-c). The key observation is high citrate, histidine and ornithine abundance in LTCS patients compared with any other group (HC, Recov, and acute) (Suppl. Fig. 6b-c). Furthermore, Spearman correlation analysis was performed to identify possible interaction among cytokines, chemokines, metabolites and lipoproteins. Herein, we identified relatively high negative correlations against 2-aminobutyrate (2-AB), an antioxidant’s synthesis controlling metabolite [56]), alanine, threonine, pyruvate, tyrosine, sarcosine, ornithine, glutamine, citrate, and several cytokine panel’s parameters (IL-10/23/12p70/8/33/6/1b/18/17A, INF-g, IFN-a2, TNF-a, and MCP-1). On the other side, the amino acid histidine was also highly elevated in the LTCS group. These results indicate a metabolic shift of LTCS individuals. Some of these findings above were confirmed in [28; 57]. As the patient’s health deteriorated, phenylalanine and histidine concentrations increased, as did ketone body levels [44]. We believe that these results are novel regarding LTCS patients.

Spearman correlation analysis with a filter of |r| ≥ 0.5 demonstrate (Fig. 4) that the acute group appeared to have a strong gender-based bias positive to (VLDL and IDL) triglycerides, succinate, 2-oxoglutaric acid (2-OG), 3-hydroxybutyrate (3-HB), acetoacetate; and a negative correlation towards HDL free cholesterol and phospholipids (Fig. 4a and Suppl. Table 7). Furthermore, several cytokine panel data entries had a positively correlation with several lipoproteins especially with VLDL triglycerides (Fig. 4a). A negative correlation among cytokines and creatinine together with sarcosine was also observed. The macrophage attractant chemokine protein MCP-1 had a special correlational profile dedicated in a r positive towards creatine and succinate whilst, r negative for the correlations with lactate, HDL free cholesterol and phospholipids. These findings highlighted a complex nexus in acute COVID-19 patients among inflammation and metabolic regulation.

**Fig. 4.**
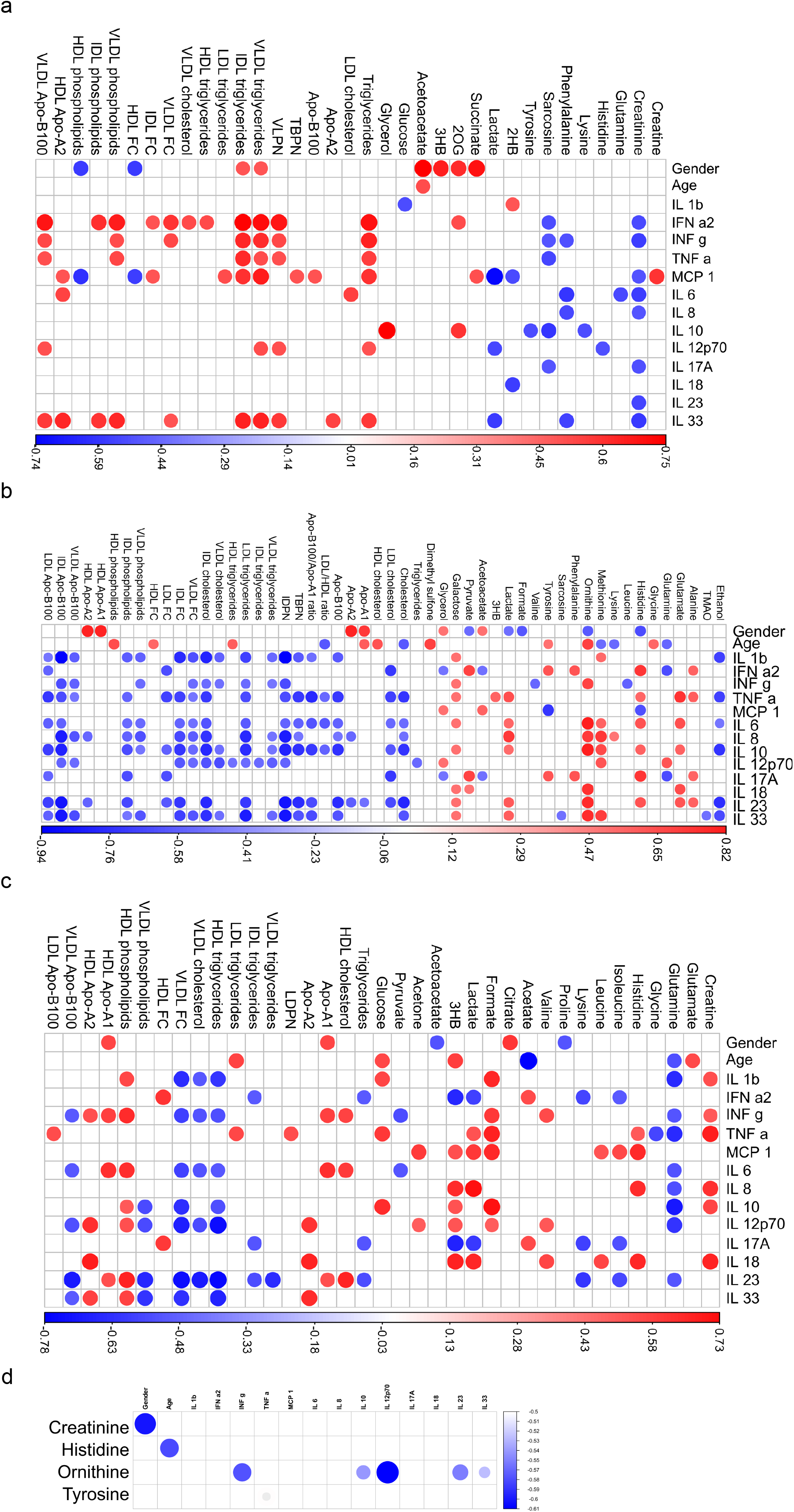
Integrated analysis of metabolites, lipoproteins, chemokines and cytokines in LTCS and comparators groups. For each patient group (Recov (EDTA plasma) n=11, HC (Heparin plasma) n=32, LTCS (EDTA plasma) n=24, acute (Heparin plasma) n=15 samples), based on the cytokine data availability, Spearman correlation test with exact p values (r values of the correlational analysis scaled −1 to 1, colored blue to red respectively) was conducted. Based on measurable metabolites data (B.I. QUANT-PS™) and a selection of lipoprotein parameter list data (B.I. LISA™). The graphical representation is performed with a filter of |r| ≥ 0.500 and has been shown in panels (sub-plot (a) – acute group, sub-plot (b) – Recov group, sub-plot (c) – LTCS group, sub-plot (d) – HC group). Correlational values for Ca-EDTA and K-EDTA not shown.

We further were interested to decipher and understand a correlation for the Recov and LTCS patients. The Recov group showed a wide spectrum of correlational dependencies, however, is based on a relatively low number of participants which must be considered as confounding factor. Nonetheless, we were able to identify a strong positive correlation among glutamate (with TNF-a), ornithine, lactate (with IL-8), and pyruvate (with IFN-a2) (Fig. 4b and Suppl. Table 8). In contrary, lipoproteins showed a mostly negative correlation to the cytokines and were mostly represented by intermediate-density lipoprotein (IDL) parameters. Most importantly, a gender-based bias positive correlation was identified for apolipoproteins A1 and A2 whilst, histidine, ornithine, and lactate negatively correlated with gender. The age appeared to be positively associated with HDL cholesterol and ornithine and negatively with overall blood cholesterol and LDL/HDL lipoprotein fractions ratio (Fig. 4b).

In case of LTCS patients some unique findings were identified. We were able to determine that a large set of cytokines were changing in a similar way to Recov amongst patients as creatine, histidine, formate, 3-HB, HDL phospholipids, HDL cholesterol, and apolipoproteins A1-A2 (Fig. 4c). Negative associations were found for glutamine, HDL triglycerides, VLDL cholesterol, and VLDL free cholesterol. Most importantly, a few gender/age-based biases were found in LTCS patients which is contrary to acute COVID-19 patients. We also noticed positive correlations of citrate and (HDL) apolipoprotein A1 with gender and negatively correlation for proline and acetoacetate. For the age-related correlations, glutamate, 3-HB, glucose, and LDL triglycerides were positively responding to the factor whilst, negative correlational response was obtained from glutamine and acetate.

In case of HC only two major negative correlations were found for the healthy controls: creatinine – gender and ornithine – IL-12p70 (Fig. 4d). Overall, it seems that each disease state has its own bubble network to combat the virus and regulate the function of host system.

## Discussion

LTCS is a condition which is thought to debilitate a person’s life after a SARS-CoV-2 viral infection and post-recovery for several months up to years. It is estimated that approximately 20-30% of all COVID-19 patients are susceptible to develop LTCS. Through our integrative metabolomics/lipoproteins and inflammation, in this finding, we identified several metabolites and lipoprotein and cytokines which are dysregulated in LTCS patient compared with either HC, Recov or acute COVID-19 patients (Fig. 5). Our major findings revealed that lactate and pyruvate were highly upregulated in LTCS patients compared with HC and similar metabolites were also upregulated in Recov patients. This could be due to dysregulated oxidative phosphorylation in Recov or LTCS patients. Furthermore, phenylalanine, glycine, acetate, Gln/Glu ratio, glutamine, and creatinine were downregulated in LTCS patients compared with HC or Recov which may be indicative of the LTCS symptom. A sign of a greater long COVID-related severity state could be demonstrated by phenylalanine, ketone bodies (acetoacetate, acetone, and 3-hydroxybutyrate), formate, creatine, and pyruvate blood levels (Fig. 5). As there is a demand for the amino acid and its further pathway products, phenylalanine levels go down in COVID recovery phase patients as reported in [58], similarly to currently investigated LTCS group versus convalescent comparison. In here, a slight change of acetoacetate could be an indicator of dietary habits changes [59] or in combination with other statistically significant parameters could predict COVID-19 disease severity [60]. From the correlational analysis, we were able to determine that correlations of creatine, glutamine, lysine, and 3-HB were stronger to the cytokine data in the LTCS group. As they had been previously reported for COVID-19-positive patients [31; 58; 61], these metabolites can provide an insight which metabolic shifts could be persisting and represent a continuous risk to the patients’ health.

**Fig. 5.**
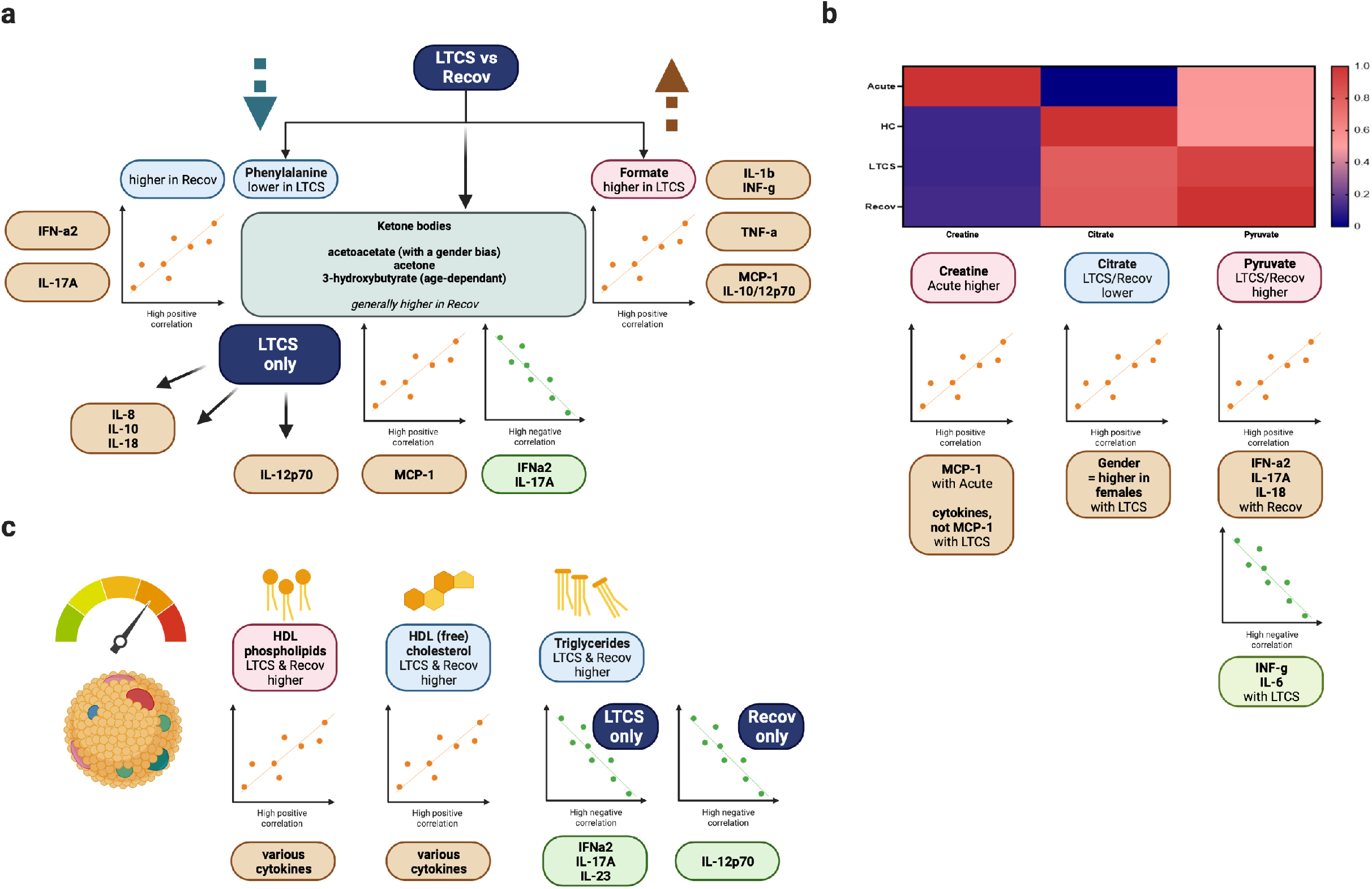
Graphical summary of the study. Sub-plot (a) focuses on phenylalanine, formate, and ketone bodies (mainly, 3-hydroxybutyrate) as significant variables identified via regression model analysis. Sub-plot (b) demonstrates further strong metabolite correlations with the cytokine data. Sub-plot (c) is showing highlights of the lipoprotein data analysis.

Recov and LTCS patients showed very similar types of metabolic dysregulations. We identified some difference between the groups especially for the formate, acetate, creatinine, HDL-4 triglycerides and HDL-4 Apo-A1 apolipoprotein levels, however no significance level was achieved. Further, we investigated a correlation between groups of Recov and LTCS. The findings therefore suggest a contrastingly higher role of creatine to IL-1b, INF-g, TNF-a (predominantly), IL-8/10/18 cytokines positive correlations among the LTCS individuals. However, our finding is similar to the previously reported association of mild/acute COVID-19 patients metabolomic analysis and its classification to the cytokine panel data [10]. Moreover, we also identified a LTCS-specific positive correlation between HDL phospholipids and IL-1b, INF-g, IL-6, IL-12p70, IL-23/33. This evidence could be complemented with finding of HDL phospholipids among other NMR-defined lipoprotein variables in COVID-19 patients [33].

We further identified an elevation of citrate and pyruvate in blood of the LTCS patient group compared with HC. This is in line with another study which identified higher levels of pyruvic acid are accumulated in the bloodstream of COVID-19 patients and which could be used to prognose disease severity [62]. Further, greater levels of lactate in COVID-19 patients are an already established finding [63]. We therefore speculate that glycolysis/gluconeogenesis and Krebs cycle metabolic pathways will lead to the elevated consumption of glucose to produce citric acid into the blood plasma. It is interesting to note that the citrate levels did not significantly correlate to any of the chemokine or cytokines as yet it was only connected to the gender factor. Therefore, gender based metabolic dysregulation could play an important role to understand the disease severity. This is especially important as certain LTCS symptoms have been reported more for female or male patients.

Maintained triglycerides and other lipoproteins changes indicate that COVID-19 like features still exist in LTCS patients when comparisons were made with HC. Elevated apolipoproteins ratio B100 to A1 and overall blood triglycerides could be attributed to the disease group [49]. Although, in our study, only triglycerides showed negative correlations to IFN-a2, IL-17A, and IL-23. IFN-is connected to the innate inflammatory reaction and this could portray the ongoing need of LTCS patients to lower down SARS-CoV-2 induced dysregulation of innate immune system [64]. Our findings are prompting to speculate that a core change in cytokine levels as well as high number of triglycerides were in present in the bloodstream of LTCS patients. This could be result of dysregulated innate immune response which could lead to a higher severity of COVID-19 like symptoms. In a similar context of lipid levels, an increase of COVID-19 severity is increased with diabetes and as a result of lowered amounts of HDL cholesterol in COVID-19 patients [65]. Our data implies that HDL cholesterol (HDCH) is lowered in the LTCS together with correlating INF-g, IL-6, and IL-23. Previously, it was identified that severe immunosuppression is key for the severity of COVID-19 rather than the cytokine storm [66]. Thus, it is plausible that lower level of lipids and inflammatory cytokines may be important for further disease symptoms in LTCS patients. Another likeliness of the LTCS group to acute COVID-19 patients is noticed *via* lowered apolipoproteins A1 & A2 levels, among other close structures they had been lowered in ill subjects [67; 68].

## Conclusion & limitation of the study

Our study provides a large set of quantitative NMR data on metabolites and lipoproteins and inflammation parameters in LTCS patients and highlights that formate, acetate, creatinine, citrate, lactate, pyruvate, histidine, ornithine, HDL and total blood triglycerides, HDL apolipoproteins Apo-A1, IL-18, TNF-a, IL-23, IL-8, MCP-1 could be key parameters in the pathophysiology of maintained disease symptoms and even progression. It should be noted as a limitation, that for this analysis only a very basic set of patient metadata is used. Thus, the estimation of the role of underlying comorbidities and the comparability to healthy controls is limited. Especially with severe COVID-19 patients (i.e. those who were hospitalized) it can be assumed that a majority of them had risk factors like diabetes, obesity, hypertension, etc. – adjustment for these risk factors in the “healthy controls” would be very interesting. The centre of this study thereof was based on the two larger groups LTCS and HC, however also within LTCS comorbidities might have contributed to changes. Furthermore, the limited number of samples for the Recov and acute group should be considered when comparing the results of this study with similar projects. Nonetheless, our results confirm and align with some of the previously published results and show novel insights into persisting altered blood metabolome, lipoproteome and inflammation parameters when comparing healthy controls with LTCS specimen.

## Authors contribution

GB: Performed the NMR spectroscopy experiments, processed the samples, analysed the integrated data, figure preparation, manuscript writing

RB: Performed the experiments, patient recruitment & processing of blood plasma AL, SG, HH, PR, MB: Patient recruitment, blood sample collection, patient metadata collection, editing the manuscript

CC, HC: control samples provided and performed analysis with IVDr (licensed)

CT: Planning of the experiments, performed the NMR spectroscopy experiments, processed the samples, analysed the integrated data, figure preparation, manuscript editing

YS: Overall project management and execution, planning of the experiments, cytokine assay and data analysis, preparation of the final figures, manuscript writing

## Acknowledgements

We thank Prof. Dr Olaf Riess for providing the infrastructure to perform the study, sharing the samples from genomic study and help with ethics writing for this project. We thank the Werner-Siemens Imaging Center with the Chair of Department Prof. Dr. Bernd Pichler for the opportunity to conduct this study. We acknowledge the Tubingen university library for open access funds for the publication.

## Data Availability Statement

Most of the data provided in the manuscript. Raw data of all the metabolite (Suppl. Table 7), lipoprotein (Suppl. Table 8), and 13-plex cytokine and age-gender data (Suppl. Table 6) are available upon request. Mean concentrations with standard deviations for each group are provided in supplementary tables 6-8.

## Figure legends

**Suppl. Fig. 1** Loadings plot of the principal component analysis (PCA, Fig. 2b).

**Suppl. Fig. 2** Identification of metabolites in LTCS and Recov patients via o PLS-DA studies. This analysis was done out based only on quantifiable metabolites data (B.I. QUANT-PS™). The 4-group distribution was shown in (a-b) using the coordinates of T score and orthogonal T score. The values that contributed the most to these VIP scores and S plot loadings of the regression model are shown here by the subplot (c). Lastly, the metabolite panel variables’ average trends were presented by sub-plot (d).

**Suppl. Fig. 3** Pathway analysis of dysregulated metabolites and lipoproteins. For each patient group pairs (Recov (EDTA plasma) n=12, HC (Heparin plasma) n=32, LTCS (EDTA plasma) n=33, acute (Heparin plasma) n=16 samples), a number of metabolic pathways (sub-plots a-f) related to the measurable metabolites data (B.I. QUANT-PS™) were identified in the patient groups. Predicted metabolic pathways are listed with p-values after the FDR applied (false discovery rate correction). The group distribution whose statistical importance has been shown in panels. Statistical significances: ns p>0.05, * p ≤0.05, ** p≤0.01, *** p≤0.001, **** p≤0.0001. 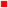 - higher average levels in a patient group; 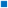 – lower average levels in a patient group. TCA cycle – tricarboxylic acid cycle. CoA – coenzyme A. ATP – adenosine triphosphate. ADP – adenosine diphosphate. SCFA – short-chain fatty acids.

**Suppl. Fig. 4** PLS-DA analysis supplement. Sub-plot is a representation of applied lipoprotein parameters in the PLS-DA analysis loadings coordinates per components 1,2 (Fig. 3a-b).

**Suppl. Fig. 5** oPLS-DA study was performed, and LTCS vs recovered patients (Recov) were compared. This analysis was done out based only on lipoprotein data panel (B.I. LISA™). The two-group distribution was shown in (a) using the coordinates of loading components 1 and 2. The values that contributed the most to these VIP scores and S-plot data of the regression model are shown here by the subplots (b) and (c), which are sorted from most significant to least significant. Lastly, the metabolite panel regression model analysis highlighted results (VIP > 1.0) visualized via a Heat map as presented by sub-plot (d).

**Suppl. Fig. 6** PLS-DA study was performed, and the available data from the cytokine panel table (Suppl. Table 6) were compared. The group distribution was shown in (a). The values that contributed the most to these loadings of the regression model are shown here by the subplots (b) by Component 1) and (c) by Component 2, which are sorted from most significant to least significant via variables in projection score (VIP) plots. Lastly, variables’ panel of Spearman correlations for IL-10 cytokines showed trends against the main cytokine parameters by sub-plot (d).

## Reference

[1] W.L. Dashboard, https://covid19.who.int. (2020).

[2] F.C. Ceballos, A. Virseda-Berdices, S. Resino, P. Ryan, O. Martinez-Gonzalez, F. Perez-Garcia, M. Martin-Vicente, O. Brochado-Kith, R. Blancas, S. Bartolome-Sanchez, E.J. Vidal-Alcantara, O.E. Alboniga-Diez, J. Cuadros-Gonzalez, N. Blanca-Lopez, I. Martinez, I.R. Martinez-Acitores, C. Barbas, A. Fernandez-Rodriguez, and M.A. Jimenez-Sousa, Metabolic Profiling at COVID-19 Onset Shows Disease Severity and Sex-Specific Dysregulation. Front Immunol 13 (2022) 925558.

[3] J.W. Lee, Y. Su, P. Baloni, D. Chen, A.J. Pavlovitch-Bedzyk, D. Yuan, V.R. Duvvuri, R.H. Ng, J. Choi, J. Xie, R. Zhang, K. Murray, S. Kornilov, B. Smith, A.T. Magis, D.S.B. Hoon, J.J. Hadlock, J.D. Goldman, N.D. Price, R. Gottardo, M.M. Davis, L. Hood, P.D. Greenberg, and J.R. Heath, Integrated analysis of plasma and single immune cells uncovers metabolic changes in individuals with COVID-19. Nat Biotechnol 40 (2022) 110–120.

[4] Y. Singh, C. Trautwein, R. Fendel, N. Krickeberg, G. Berezhnoy, R. Bissinger, S. Ossowski, M.S. Salker, N. Casadei, O. Riess, and C.-O.I. Deutsche, SARS-CoV-2 infection paralyzes cytotoxic and metabolic functions of the immune cells. Heliyon 7 (2021) e07147.

[5] X. Liu, J. Zhao, H. Wang, W. Wang, X. Su, X. Liao, S. Zhang, J. Sun, and Z. Zhang, Metabolic Defects of Peripheral T Cells in COVID-19 Patients. J Immunol 206 (2021) 2900–2908.

[6] S.M. O’Carroll, and L.A.J. O’Neill, Targeting immunometabolism to treat COVID-19. Immunother Adv 1 (2021) ltab013.

[7] M. Cornillet, B. Strunz, O. Rooyackers, A. Ponzetta, P. Chen, J.R. Muvva, M. Akber, M. Buggert, B.J. Chambers, M. Dzidic, I. Filipovic, J.B. Gorin, S. Gredmark-Russ, L. Hertwig, J. Klingstrom, E. Kokkinou, E. Kvedaraite, M. Lourda, J. Mjosberg, C. Maucourant, A. Norrby-Teglund, T. Parrot, A. Perez-Potti, O. Rivera-Ballesteros, J.K. Sandberg, J.T. Sandberg, T. Sekine, M. Svensson, R. Varnaite, K.I.K.C.-S.G. Karolinska, L.I. Eriksson, S. Aleman, K. Stralin, H.G. Ljunggren, and N.K. Bjorkstrom, COVID-19- specific metabolic imprint yields insights into multiorgan system perturbations. Eur J Immunol 52 (2022) 503–510.

[8] H. Jia, C. Liu, D. Li, Q. Huang, D. Liu, Y. Zhang, C. Ye, D. Zhou, Y. Wang, Y. Tan, K. Li, F. Lin, H. Zhang, J. Lin, Y. Xu, J. Liu, Q. Zeng, J. Hong, G. Chen, H. Zhang, L. Zheng, X. Deng, C. Ke, Y. Gao, J. Fan, B. Di, and H. Liang, Metabolomic analyses reveal new stagespecific features of COVID-19. Eur Respir J 59 (2022).

[9] M. Costanzo, M. Caterino, R. Fedele, A. Cevenini, M. Pontillo, L. Barra, and M. Ruoppolo, COVIDomics: The Proteomic and Metabolomic Signatures of COVID-19. Int J Mol Sci 23 (2022).

[10] F.X. Danlos, C. Grajeda-Iglesias, S. Durand, A. Sauvat, M. Roumier, D. Cantin, E. Colomba, J. Rohmer, F. Pommeret, G. Baciarello, C. Willekens, M. Vasse, F. Griscelli, J.E. Fahrner, A.G. Goubet, A. Dubuisson, L. Derosa, N. Nirmalathasan, D. Bredel, S. Mouraud, C. Pradon, A. Stoclin, F. Rozenberg, J. Duchemin, G. Jourdi, S. Ellouze, F. Levavasseur, L. Albiges, J.C. Soria, F. Barlesi, E. Solary, F. Andre, F. Pene, F. Ackerman, L. Mouthon, L. Zitvogel, A. Marabelle, J.M. Michot, M. Fontenay, and G. Kroemer, Metabolomic analyses of COVID-19 patients unravel stage-dependent and prognostic biomarkers. Cell Death Dis 12 (2021) 258.

[11] T.S. Di Wu, Xiaobo Yang, Jian-Xin Song Mingliang Zhang Chengye Yao Wen Liu, Muhan Huang, Yuan Yu, Qingyu Yang Tingju Zhu, Jiqian Xu, Jingfang Mu, Yaxin Wang, Hong Wang, Tang Tang, Yujie Ren Yongran Wu, Shu-Hai Lin, Yang Qiu, Ding-Yu Zhang, You Shang, Xi Zhou, Plasma Metabolomic and Lipidomic Alterations Associated with COVID-19. Natl Sci Rev (2020).

[12] S. Lodge, P. Nitschke, T. Kimhofer, J.D. Coudert, S. Begum, S.H. Bong, T. Richards, D. Edgar, E. Raby, M. Spraul, H. Schaefer, J.C. Lindon, R.L. Loo, E. Holmes, and J.K. Nicholson, NMR Spectroscopic Windows on the Systemic Effects of SARS-CoV-2 Infection on Plasma Lipoproteins and Metabolites in Relation to Circulating Cytokines. J Proteome Res 20 (2021) 1382–1396.

[13] Q. Wan, M. Chen, Z. Zhang, Y. Yuan, H. Wang, Y. Hao, W. Nie, L. Wu, and S. Chen, Machine Learning of Serum Metabolic Patterns Encodes Asymptomatic SARS-CoV-2 Infection. Front Chem 9 (2021) 746134.

[14] M.A. Hassan, K. Al-Sakkaf, M.R. Shait Mohammed, A. Dallol, J. Al-Maghrabi, A. Aldahlawi, S. Ashoor, M. Maamra, J. Ragoussis, W. Wu, M.I. Khan, A.L. Al-Malki, and H. Choudhry, Integration of Transcriptome and Metabolome Provides Unique Insights to Pathways Associated With Obese Breast Cancer Patients. Front Oncol 10 (2020)804.

[15] Y. Su, D. Chen, D. Yuan, C. Lausted, J. Choi, C.L. Dai, V. Voillet, V.R. Duvvuri, K. Scherler, P. Troisch, P. Baloni, G. Qin, B. Smith, S.A. Kornilov, C. Rostomily, A. Xu, J. Li, S. Dong, A. Rothchild, J. Zhou, K. Murray, R. Edmark, S. Hong, J.E. Heath, J. Earls, R. Zhang, J. Xie, S. Li, R. Roper, L. Jones, Y. Zhou, L. Rowen, R. Liu, S. Mackay, D.S. O’Mahony, C.R. Dale, J.A. Wallick, H.A. Algren, M.A. Zager, I.S.-S.C.B. Unit, W. Wei, N.D. Price, S. Huang, N. Subramanian, K. Wang, A.T. Magis, J.J. Hadlock, L. Hood, A. Aderem, J.A. Bluestone, L.L. Lanier, P.D. Greenberg, R. Gottardo, M.M. Davis, J.D. Goldman, and J.R. Heath, Multi-Omics Resolves a Sharp Disease-State Shift between Mild and Moderate COVID-19. Cell 183 (2020) 1479–1495 e20.

[16] F. He, T. Zhang, K. Xue, Z. Fang, G. Jiang, S. Huang, K. Li, Z. Gu, H. Shi, Z. Zhang, H. Zhu, L. Lin, J. Li, F. Xiao, H. Shan, R. Yan, X. Li, and Z. Yan, Fecal multi-omics analysis reveals diverse molecular alterations of gut ecosystem in COVID-19 patients. Anal Chim Acta 1180 (2021)338881.

[17] L. Lv, H. Jiang, Y. Chen, S. Gu, J. Xia, H. Zhang, Y. Lu, R. Yan, and L. Li, The faecal metabolome in COVID-19 patients is altered and associated with clinical features and gut microbes. Anal Chim Acta 1152 (2021) 338267.

[18] C. Wang, X. Li, W. Ning, S. Gong, F. Yang, C. Fang, Y. Gong, D. Wu, M. Huang, Y. Gou, S. Fu, Y. Ren, R. Yang, Y. Qiu, Y. Xue, Y. Xu, and X. Zhou, Multi-omic profiling of plasma reveals molecular alterations in children with COVID-19. Theranostics 11 (2021) 8008–8026.

[19] P. Wu, D. Chen, W. Ding, P. Wu, H. Hou, Y. Bai, Y. Zhou, K. Li, S. Xiang, P. Liu, J. Ju, E. Guo, J. Liu, B. Yang, J. Fan, L. He, Z. Sun, L. Feng, J. Wang, T. Wu, H. Wang, J. Cheng, H. Xing, Y. Meng, Y. Li, Y. Zhang, H. Luo, G. Xie, X. Lan, Y. Tao, J. Li, H. Yuan, K. Huang, W. Sun, X. Qian, Z. Li, M. Huang, P. Ding, H. Wang, J. Qiu, F. Wang, S. Wang, J. Zhu, X. Ding, C. Chai, L. Liang, X. Wang, L. Luo, Y. Sun, Y. Yang, Z. Zhuang, T. Li, L. Tian, S. Zhang, L. Zhu, A. Chang, L. Chen, Y. Wu, X. Ma, F. Chen, Y. Ren, X. Xu, S. Liu, J. Wang, H. Yang, L. Wang, C. Sun, D. Ma, X. Jin, and G. Chen, The trans-omics landscape of COVID-19. Nat Commun 12 (2021) 4543.

[20] Z. Song, L. Bao, W. Deng, J. Liu, E. Ren, Q. Lv, M. Liu, F. Qi, T. Chen, R. Deng, F. Li, Y. Liu, Q. Wei, H. Gao, P. Yu, Y. Han, W. Zhao, J. Zheng, X. Liang, F. Yang, and C. Qin, Integrated histopathological, lipidomic, and metabolomic profiles reveal mink is a useful animal model to mimic the pathogenicity of severe COVID-19 patients. Signal Transduct Target Ther 7 (2022) 29.

[21] Y.M. Chen, Y. Zheng, Y. Yu, Y. Wang, Q. Huang, F. Qian, L. Sun, Z.G. Song, Z. Chen, J. Feng, Y. An, J. Yang, Z. Su, S. Sun, F. Dai, Q. Chen, Q. Lu, P. Li, Y. Ling, Z. Yang, H. Tang, L. Shi, L. Jin, E.C. Holmes, C. Ding, T.Y. Zhu, and Y.Z. Zhang, Blood molecular markers associated with COVID-19 immunopathology and multi-organ damage. EMBO J 39 (2020) e105896.

[22] M. Bizkarguenaga, C. Bruzzone, R. Gil-Redondo, I. SanJuan, I. Martin-Ruiz, D. Barriales, A. Palacios, S.T. Pasco, B. Gonzalez-Valle, A. Lain, L. Herrera, A. Azkarate, M.A. Vesga, C. Eguizabal, J. Anguita, N. Embade, J.M. Mato, and O. Millet, Uneven metabolic and lipidomic profiles in recovered COVID-19 patients as investigated by plasma NMR metabolomics. NMR Biomed 35 (2022) e4637.

[23] S. Falck-Jones, S. Vangeti, M. Yu, R. Falck-Jones, A. Cagigi, I. Badolati, B. Osterberg, M.J. Lautenbach, E. Ahlberg, A. Lin, R. Lepzien, I. Szurgot, K. Lenart, F. Hellgren, H. Maecker, J. Salde, J. Albert, N. Johansson, M. Bell, K. Lore, A. Farnert, and A. Smed-Sorensen, Functional monocytic myeloid-derived suppressor cells increase in blood but not airways and predict COVID-19 severity. J Clin Invest 131 (2021).

[24] M.S. Abers, O.M. Delmonte, E.E. Ricotta, J. Fintzi, D.L. Fink, A.A.A. de Jesus, K.A. Zarember, S. Alehashemi, V. Oikonomou, J.V. Desai, S.W. Canna, B. Shakoory, K. Dobbs, L. Imberti, A. Sottini, E. Quiros-Roldan, F. Castelli, C. Rossi, D. Brugnoni, A. Biondi, L.R. Bettini, M. D’Angio, P. Bonfanti, R. Castagnoli, D. Montagna, A. Licari, G.L. Marseglia, E.F. Gliniewicz, E. Shaw, D.E. Kahle, A.T. Rastegar, M. Stack, K. Myint-Hpu, S.L. Levinson, M.J. DiNubile, D.W. Chertow, P.D. Burbelo, J.I. Cohen, K.R. Calvo, J.S. Tsang, N.C.-. Consortium, H.C. Su, J.I. Gallin, D.B. Kuhns, R. Goldbach-Mansky, M.S. Lionakis, and L.D. Notarangelo, An immune-based biomarker signature is associated with mortality in COVID-19 patients. JCI Insight 6 (2021).

[25] Z. Al-Aly, B. Bowe, and Y. Xie, Long COVID after breakthrough SARS-CoV-2 infection. Nat Med 28 (2022) 1461–1467.

[26] V.P. Guntur, T. Nemkov, E. de Boer, M.P. Mohning, D. Baraghoshi, F.I. Cendali, I. San-Millan, I. Petrache, and A. D’Alessandro, Signatures of Mitochondrial Dysfunction and Impaired Fatty Acid Metabolism in Plasma of Patients with Post-Acute Sequelae of COVID-19 (PASC). Metabolites 12 (2022).

[27] H.L. Correa, L.A. Deus, T.B. Araujo, A.L. Reis, C.E.N. Amorim, A.B. Gadelha, R.L. Santos, F.S. Honorato, D. Motta-Santos, C. Tzanno-Martins, R.V.P. Neves, and T.S. Rosa, Phosphate and IL-10 concentration as predictors of long-covid in hemodialysis patients: A Brazilian study. Front Immunol 13 (2022) 1006076.

[28] T. Kimhofer, S. Lodge, L. Whiley, N. Gray, R.L. Loo, N.G. Lawler, P. Nitschke, S.-H. Bong, D.L. Morrison, S. Begum, T. Richards, B.B. Yeap, C. Smith, K.G.C. Smith, E. Holmes, and J.K. Nicholson, Integrative Modeling of Quantitative Plasma Lipoprotein, Metabolic, and Amino Acid Data Reveals a Multiorgan Pathological Signature of SARS-CoV-2 Infection. J. Proteome Res. 19 (2020) 4442–4454.

[29] B.S.B. Correia, V.G. Ferreira, P.M.F.D. Piagge, M.B. Almeida, N.A. Assunção, J.R.S. Raimundo, F.L.A. Fonseca, E. Carrilho, and D.R. Cardoso, 1H qNMR-Based Metabolomics Discrimination of Covid-19 Severity. J. Proteome Res. 21 (2022) 1640–1653.

[30] V. Ghini, L. Maggi, A. Mazzoni, M. Spinicci, L. Zammarchi, A. Bartoloni, F. Annunziato, and P. Turano, Serum NMR Profiling Reveals Differential Alterations in the Lipoproteome Induced by Pfizer-BioNTech Vaccine in COVID-19 Recovered Subjects and Naïve Subjects. Frontiers in molecular biosciences 9 (2022) 839809.

[31] G. Meoni, V. Ghini, L. Maggi, A. Vignoli, A. Mazzoni, L. Salvati, M. Capone, A. Vanni, L. Tenori, P. Fontanari, F. Lavorini, A. Peris, A. Bartoloni, F. Liotta, L. Cosmi, C. Luchinat, F. Annunziato, and P. Turano, Metabolomic/lipidomic profiling of COVID-19 and individual response to tocilizumab. PLoS Pathog 17 (2021) e1009243.

[32] E. Holmes, J. Wist, R. Masuda, S. Lodge, P. Nitschke, T. Kimhofer, R.L. Loo, S. Begum, B. Boughton, R. Yang, A.-C. Morillon, S.-T. Chin, D. Hall, M. Ryan, S.-H. Bong, M. Gay, D.W. Edgar, J.C. Lindon, T. Richards, B.B. Yeap, S. Pettersson, M. Spraul, H. Schaefer, N.G. Lawler, N. Gray, L. Whiley, and J.K. Nicholson, Incomplete Systemic Recovery and Metabolic Phenoreversion in Post-Acute-Phase Nonhospitalized COVID-19 Patients: Implications for Assessment of Post-Acute COVID-19 Syndrome. J. Proteome Res. 20 (2021) 3315–3329.

[33] R. Masuda, S. Lodge, P. Nitschke, M. Spraul, H. Schaefer, S.H. Bong, T. Kimhofer, D. Hall, R.L. Loo, M. Bizkarguenaga, C. Bruzzone, R. Gil-Redondo, N. Embade, J.M. Mato, E. Holmes, J. Wist, O. Millet, and J.K. Nicholson, Integrative Modeling of Plasma Metabolic and Lipoprotein Biomarkers of SARS-CoV-2 Infection in Spanish and Australian COVID-19 Patient Cohorts. J Proteome Res 20 (2021) 4139–4152.

[34] P. Nitschke, S. Lodge, D. Hall, H. Schaefer, M. Spraul, N. Embade, O. Millet, E. Holmes, J. Wist, and J.K. Nicholson, Direct low field J-edited diffusional proton NMR spectroscopic measurement of COVID-19 inflammatory biomarkers in human serum. Analyst 147 (2022) 4213–4221.

[35] F. Schmelter, B. Föh, A. Mallagaray, J. Rahmöller, M. Ehlers, S. Lehrian, V. von Kopylow, I. Künsting, A.S. Lixenfeld, E. Martin, M. Ragab, R. Meyer-Saraei, F. Kreutzmann, I. Eitel, S. Taube, N. Käding, E. Jantzen, T. Graf, C. Sina, and U.L. Günther, Metabolic and Lipidomic Markers Differentiate COVID-19 From Non-Hospitalized and Other Intensive Care Patients. 8 (2021).

[36] T. Rossler, G. Berezhnoy, Y. Singh, C. Cannet, T. Reinsperger, H. Schafer, M. Spraul, M. Kneilling, U. Merle, and C. Trautwein, Quantitative Serum NMR Spectroscopy Stratifies COVID-19 Patients and Sheds Light on Interfaces of Host Metabolism and the Immune Response with Cytokines and Clinical Parameters. Metabolites 12 (2022).

[37] W. Stacklies, H. Redestig, M. Scholz, D. Walther, and J. Selbig, pcaMethods--a bioconductor package providing PCA methods for incomplete data. Bioinformatics 23 (2007) 1164–7.

[38] F. Dieterle, A. Ross, G. Schlotterbeck, and H. Senn, Probabilistic quotient normalization as robust method to account for dilution of complex biological mixtures. Application in 1H NMR metabonomics. Anal Chem 78 (2006) 4281–90.

[39] Z. Pang, G. Zhou, J. Chong, and J. Xia, Comprehensive Meta-Analysis of COVID-19 Global Metabolomics Datasets. Metabolites 11 (2021).

[40] N. Karu, A. Kindt, A.J. van Gammeren, A.A.M. Ermens, A.C. Harms, L. Portengen, R.C.H. Vermeulen, W.A. Dik, A.W. Langerak, V.H.J. van der Velden, and T. Hankemeier, Severe COVID-19 Is Characterised by Perturbations in Plasma Amines Correlated with Immune Response Markers, and Linked to Inflammation and Oxidative Stress. Metabolites 12 (2022).

[41] G. Kaur, X. Ji, and I. Rahman, SARS-CoV2 Infection Alters Tryptophan Catabolism and Phospholipid Metabolism. Metabolites 11 (2021).

[42] S. Krishnan, H. Nordqvist, A.T. Ambikan, S. Gupta, M. Sperk, S. Svensson-Akusjarvi, F. Mikaeloff, R. Benfeitas, E. Saccon, S.M. Ponnan, J.E. Rodriguez, N. Nikouyan, A. Odeh, G. Ahlen, M. Asghar, M. Sallberg, J. Vesterbacka, P. Nowak, A. Vegvari, A. Sonnerborg, C.J. Treutiger, and U. Neogi, Metabolic Perturbation Associated With COVID-19 Disease Severity and SARS-CoV-2 Replication. Mol Cell Proteomics 20 (2021) 100159.

[43] H.M. Lewis, Y. Liu, C.F. Frampas, K. Longman, M. Spick, A. Stewart, E. Sinclair, N. Kasar, D. Greener, A.D. Whetton, P.E. Barran, T. Chen, D. Dunn-Walters, D.J. Skene, and M.J. Bailey, Metabolomics Markers of COVID-19 Are Dependent on Collection Wave. Metabolites 12 (2022).

[44] C. Bruzzone, M. Bizkarguenaga, R. Gil-Redondo, T. Diercks, E. Arana, A. Garcia de Vicuna, M. Seco, A. Bosch, A. Palazon, I. San Juan, A. Lain, J. Gil-Martinez, G. Bernardo-Seisdedos, D. Fernandez-Ramos, F. Lopitz-Otsoa, N. Embade, S. Lu, J.M. Mato, and O. Millet, SARS-CoV-2 Infection Dysregulates the Metabolomic and Lipidomic Profiles of Serum. iScience 23 (2020) 101645.

[45] B.S.B. Correia, V.G. Ferreira, P. Piagge, M.B. Almeida, N.A. Assuncao, J.R.S. Raimundo, F.L.A. Fonseca, E. Carrilho, and D.R. Cardoso, (1)H qNMR-Based Metabolomics Discrimination of Covid-19 Severity. J Proteome Res 21 (2022) 1640–1653.

[46] A.R. Gafson, T. Thorne, C.I.J. McKechnie, B. Jimenez, R. Nicholas, and P.M. Matthews, Lipoprotein markers associated with disability from multiple sclerosis. Sci Rep 8 (2018) 17026.

[47] M. Sindelar, E. Stancliffe, M. Schwaiger-Haber, D.S. Anbukumar, K. Adkins-Travis, C.W. Goss, J.A. O’Halloran, P.A. Mudd, W.C. Liu, R.A. Albrecht, A. Garcia-Sastre, L.P. Shriver, and G.J. Patti, Longitudinal metabolomics of human plasma reveals prognostic markers of COVID-19 disease severity. Cell Rep Med 2 (2021) 100369.

[48] C. Bruzzone, M. Bizkarguenaga, R. Gil-Redondo, T. Diercks, E. Arana, A. García de Vicuña, M. Seco, A. Bosch, A. Palazón, I. San Juan, A. Laín, J. Gil-Martínez, G. Bernardo-Seisdedos, D. Fernández-Ramos, F. Lopitz-Otsoa, N. Embade, S. Lu, J.M. Mato, and O. Millet, SARS-CoV-2 Infection Dysregulates the Metabolomic and Lipidomic Profiles of Serum. iScience 23 (2020) 101645.

[49] S. Lodge, P. Nitschke, T. Kimhofer, J.D. Coudert, S. Begum, S.-H. Bong, T. Richards, D. Edgar, E. Raby, M. Spraul, H. Schaefer, J.C. Lindon, R.L. Loo, E. Holmes, and J.K. Nicholson, NMR Spectroscopic Windows on the Systemic Effects of SARS-CoV-2 Infection on Plasma Lipoproteins and Metabolites in Relation to Circulating Cytokines. J. Proteome Res. 20 (2021) 1382–1396.

[50] M. Buyukozkan, S. Alvarez-Mulett, A.C. Racanelli, F. Schmidt, R. Batra, K.L. Hoffman, H. Sarwath, R. Engelke, L. Gomez-Escobar, W. Simmons, E. Benedetti, K. Chetnik, G. Zhang, E. Schenck, K. Suhre, J.J. Choi, Z. Zhao, S. Racine-Brzostek, H.S. Yang, M.E. Choi, A.M.K. Choi, S.J. Cho, and J. Krumsiek, Integrative metabolomic and proteomic signatures define clinical outcomes in severe COVID-19. iScience 25 (2022) 104612.

[51] V. Ghini, G. Meoni, L. Pelagatti, T. Celli, F. Veneziani, F. Petrucci, V. Vannucchi, L. Bertini, C. Luchinat, G. Landini, and P. Turano, Profiling metabolites and lipoproteins in COMETA, an Italian cohort of COVID-19 patients. PLoS Pathog 18 (2022) e1010443.

[52] J.J. Kovarik, A. Bileck, G. Hagn, S.M. Meier-Menches, T. Frey, A. Kaempf, M. Hollenstein, T. Shoumariyeh, L. Skos, B. Reiter, M.C. Gerner, A. Spannbauer, E. Hasimbegovic, D. Schmidl, G. Garhöfer, M. Gyöngyösi, K.G. Schmetterer, and C. Gerner, (2022).

[53] P.E. Scherer, J.P. Kirwan, and C.J. Rosen, Post-acute sequelae of COVID-19: A metabolic perspective. Elife 11 (2022).

[54] J. Liu, S. Li, J. Liu, B. Liang, X. Wang, H. Wang, W. Li, Q. Tong, J. Yi, L. Zhao, L. Xiong, C. Guo, J. Tian, J. Luo, J. Yao, R. Pang, H. Shen, C. Peng, T. Liu, Q. Zhang, J. Wu, L. Xu, S. Lu, B. Wang, Z. Weng, C. Han, H. Zhu, R. Zhou, H. Zhou, X. Chen, P. Ye, B. Zhu, L. Wang, W. Zhou, S. He, Y. He, S. Jie, P. Wei, J. Zhang, Y. Lu, W. Wang, L. Zhang, L. Li, F. Zhou, J. Wang, U. Dittmer, M. Lu, Y. Hu, D. Yang, and X. Zheng, Longitudinal characteristics of lymphocyte responses and cytokine profiles in the peripheral blood of SARS-CoV-2 infected patients. EBioMedicine 55 (2020) 102763.

[55] D.M. Del Valle, S. Kim-Schulze, H.-H. Huang, N.D. Beckmann, S. Nirenberg, B. Wang, Y. Lavin, T.H. Swartz, D. Madduri, A. Stock, T.U. Marron, H. Xie, M. Patel, K. Tuballes, O. Van Oekelen, A. Rahman, P. Kovatch, J.A. Aberg, E. Schadt, S. Jagannath, M. Mazumdar, A.W. Charney, A. Firpo-Betancourt, D.R. Mendu, J. Jhang, D. Reich, K. Sigel, C. Cordon-Cardo, M. Feldmann, S. Parekh, M. Merad, and S. Gnjatic, An inflammatory cytokine signature predicts COVID-19 severity and survival. Nature Medicine (2020).

[56] Y. Irino, R. Toh, M. Nagao, T. Mori, T. Honjo, M. Shinohara, S. Tsuda, H. Nakajima, S. Satomi-Kobayashi, T. Shinke, H. Tanaka, T. Ishida, O. Miyata, and K.-i. Hirata, 2- Aminobutyric acid modulates glutathione homeostasis in the myocardium. Sci Rep 6 (2016)36749.

[57] D. Shi, R. Yan, L. Lv, H. Jiang, Y. Lu, J. Sheng, J. Xie, W. Wu, J. Xia, K. Xu, S. Gu, Y. Chen, C. Huang, J. Guo, Y. Du, and L. Li, The serum metabolome of COVID-19 patients is distinctive and predictive. Metabolism 118 (2021) 154739.

[58] L. Ansone, M. Briviba, I. Silamikelis, A. Terentjeva, I. Perkons, L. Birzniece, V. Rovite, B. Rozentale, L. Viksna, O. Kolesova, K. Klavins, and J. Klovins, Amino Acid Metabolism is Significantly Altered at the Time of Admission in Hospital for Severe COVID-19 Patients: Findings from Longitudinal Targeted Metabolomics Analysis. 9 (2021) e00338–21.

[59] M. Watanabe, A. Balena, D. Masi, R. Tozzi, R. Risi, A. Caputi, R. Rossetti, M.E. Spoltore, F. Biagi, E. Anastasi, A. Angeloni, S. Mariani, C. Lubrano, D. Tuccinardi, and L. Gnessi, Rapid Weight Loss, Central Obesity Improvement and Blood Glucose Reduction Are Associated with a Stronger Adaptive Immune Response Following COVID-19 mRNA Vaccine. 10 (2022)79.

[60] J.W. Lee, Y. Su, P. Baloni, D. Chen, A.J. Pavlovitch-Bedzyk, D. Yuan, V.R. Duvvuri, R.H. Ng, J. Choi, J. Xie, R. Zhang, K. Murray, S. Kornilov, B. Smith, A.T. Magis, D.S.B. Hoon, J.J. Hadlock, J.D. Goldman, N.D. Price, R. Gottardo, M.M. Davis, L. Hood, P.D. Greenberg, and J.R. Heath, Integrated analysis of plasma and single immune cells uncovers metabolic changes in individuals with COVID-19. Nature Biotechnology 40 (2022) 110–120.

[61] Y. Su, D. Chen, D. Yuan, C. Lausted, J. Choi, C.L. Dai, V. Voillet, V.R. Duvvuri, K. Scherler, P. Troisch, P. Baloni, G. Qin, B. Smith, S.A. Kornilov, C. Rostomily, A. Xu, J. Li, S. Dong, A. Rothchild, J. Zhou, K. Murray, R. Edmark, S. Hong, J.E. Heath, J. Earls, R. Zhang, J. Xie, S. Li, R. Roper, L. Jones, Y. Zhou, L. Rowen, R. Liu, S. Mackay, D.S. O’Mahony, C.R. Dale, J.A. Wallick, H.A. Algren, M.A. Zager, W. Wei, N.D. Price, S. Huang, N. Subramanian, K. Wang, A.T. Magis, J.J. Hadlock, L. Hood, A. Aderem, J.A. Bluestone, L.L. Lanier, P.D. Greenberg, R. Gottardo, M.M. Davis, J.D. Goldman, and J.R. Heath, Multi-Omics Resolves a Sharp Disease-State Shift between Mild and Moderate COVID-19. Cell 183 (2020) 1479–1495.e20.

[62] V. Ceperuelo-Mallafre, L. Reverte, J. Peraire, A. Madeira, E. Maymo-Masip, M. Lopez-Dupla, A. Gutierrez-Valencia, E. Ruiz-Mateos, M.J. Buzon, R. Jorba, J. Vendrell, T. Auguet, M. Olona, F. Vidal, A. Rull, and S. Fernandez-Veledo, Circulating pyruvate is a potent prognostic marker for critical COVID-19 outcomes. Front Immunol 13 (2022) 912579.

[63] G. Carpenè, D. Onorato, R. Nocini, G. Fortunato, J.G. Rizk, B.M. Henry, and G. Lippi, Blood lactate concentration in COVID-19: a systematic literature review. 60 (2022) 332–337.

[64] Z. Wang, H. Pan, and B. Jiang, Type I IFN deficiency: an immunological characteristic of severe COVID-19 patients. Signal Transduct Target Ther 5 (2020) 198.

[65] G. Targher, A. Mantovani, X.B. Wang, H.D. Yan, Q.F. Sun, K.H. Pan, C.D. Byrne, K.I. Zheng, Y.P. Chen, M. Eslam, J. George, and M.H. Zheng, Patients with diabetes are at higher risk for severe illness from COVID-19. Diabetes Metab 46 (2020) 335–337.

[66] K.E. Remy, M. Mazer, D.A. Striker, A.H. Ellebedy, A.H. Walton, J. Unsinger, T.M. Blood, P.A. Mudd, D.J. Yi, D.A. Mannion, D.F. Osborne, R.S. Martin, N.J. Anand, J.P. Bosanquet, J. Blood, A.M. Drewry, C.C. Caldwell, I.R. Turnbull, S.C. Brakenridge, L.L. Moldwawer, and R.S. Hotchkiss, Severe immunosuppression and not a cytokine storm characterizes COVID-19 infections. JCI Insight 5 (2020).

[67] C.B. Messner, V. Demichev, D. Wendisch, L. Michalick, M. White, A. Freiwald, K. Textoris-Taube, S.I. Vernardis, A.S. Egger, M. Kreidl, D. Ludwig, C. Kilian, F. Agostini, A. Zelezniak, C. Thibeault, M. Pfeiffer, S. Hippenstiel, A. Hocke, C. von Kalle, A. Campbell, C. Hayward, D.J. Porteous, R.E. Marioni, C. Langenberg, K.S. Lilley, W.M. Kuebler, M. Mulleder, C. Drosten, N. Suttorp, M. Witzenrath, F. Kurth, L.E. Sander, and M. Ralser, Ultra-High-Throughput Clinical Proteomics Reveals Classifiers of COVID-19 Infection. Cell Syst 11 (2020) 11–24 e4.

[68] B. Shen, X. Yi, Y. Sun, X. Bi, J. Du, C. Zhang, S. Quan, F. Zhang, R. Sun, L. Qian, W. Ge, W. Liu, S. Liang, H. Chen, Y. Zhang, J. Li, J. Xu, Z. He, B. Chen, J. Wang, H. Yan, Y. Zheng, D. Wang, J. Zhu, Z. Kong, Z. Kang, X. Liang, X. Ding, G. Ruan, N. Xiang, X. Cai, H. Gao, L. Li, S. Li, Q. Xiao, T. Lu, Y. Zhu, H. Liu, H. Chen, and T. Guo, Proteomic and Metabolomic Characterization of COVID-19 Patient Sera. Cell 182 (2020) 59–72 e15.

