## Supplementary figures and images for "Maintained imbalance of triglycerides, apolipoproteins, energy metabolites and cytokines in long-term COVID-19 syndrome (LTCS) patients"

### Suppl Fig

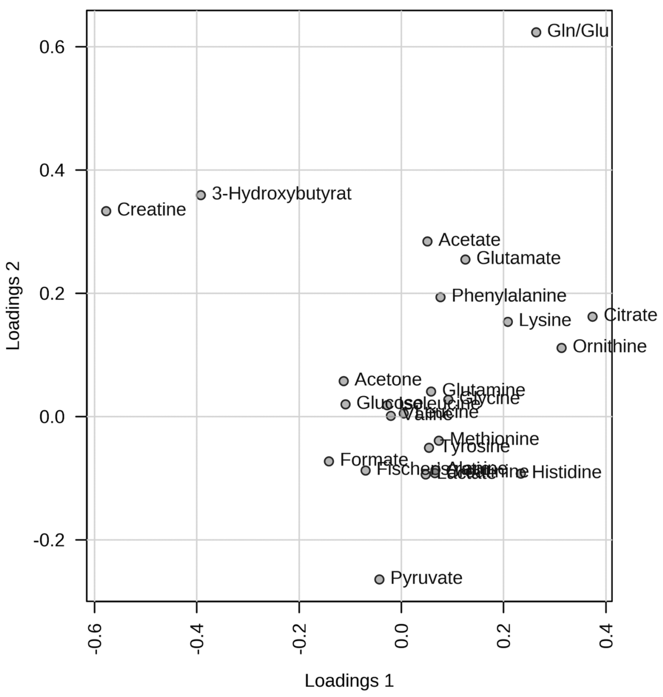

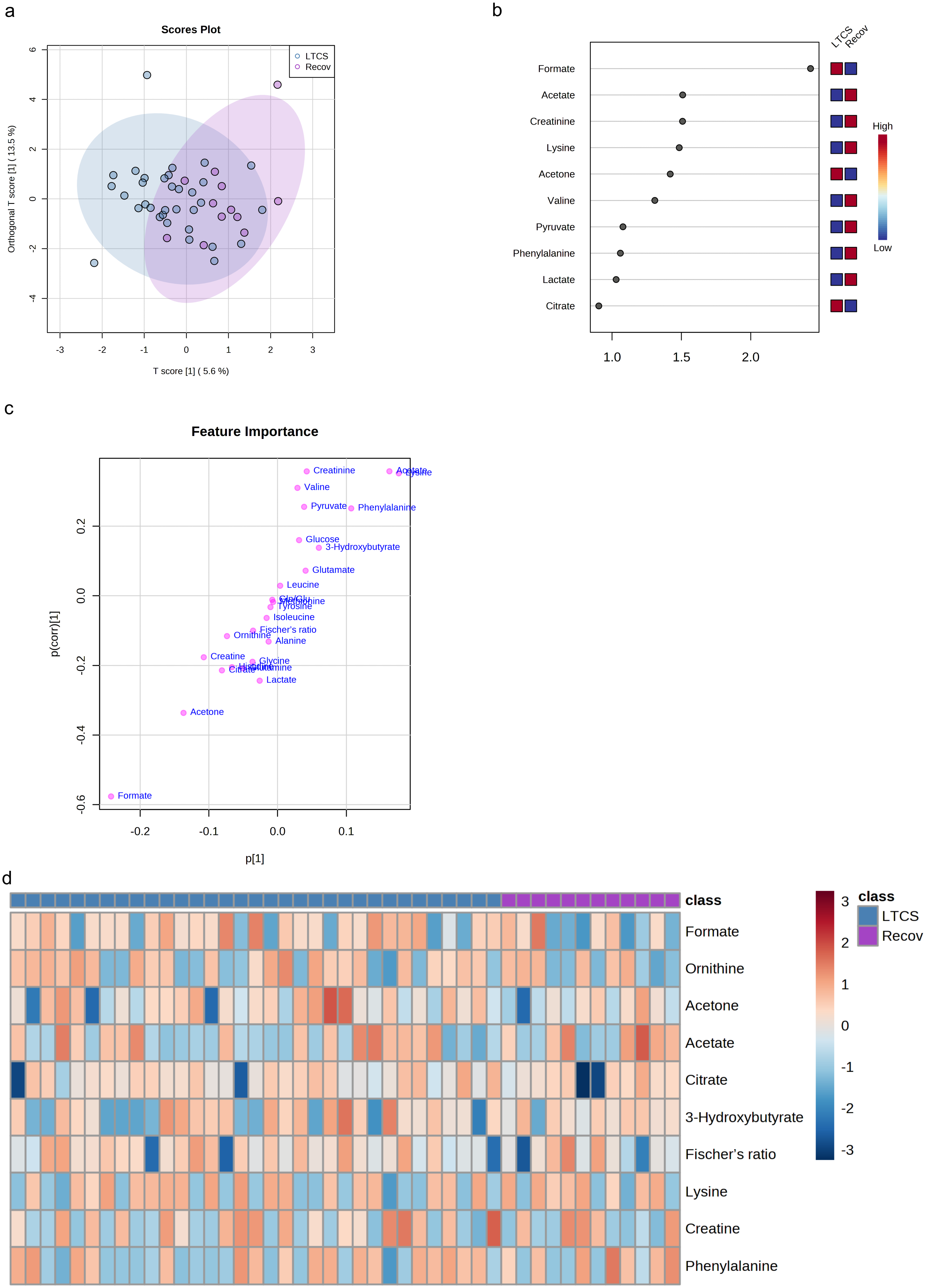

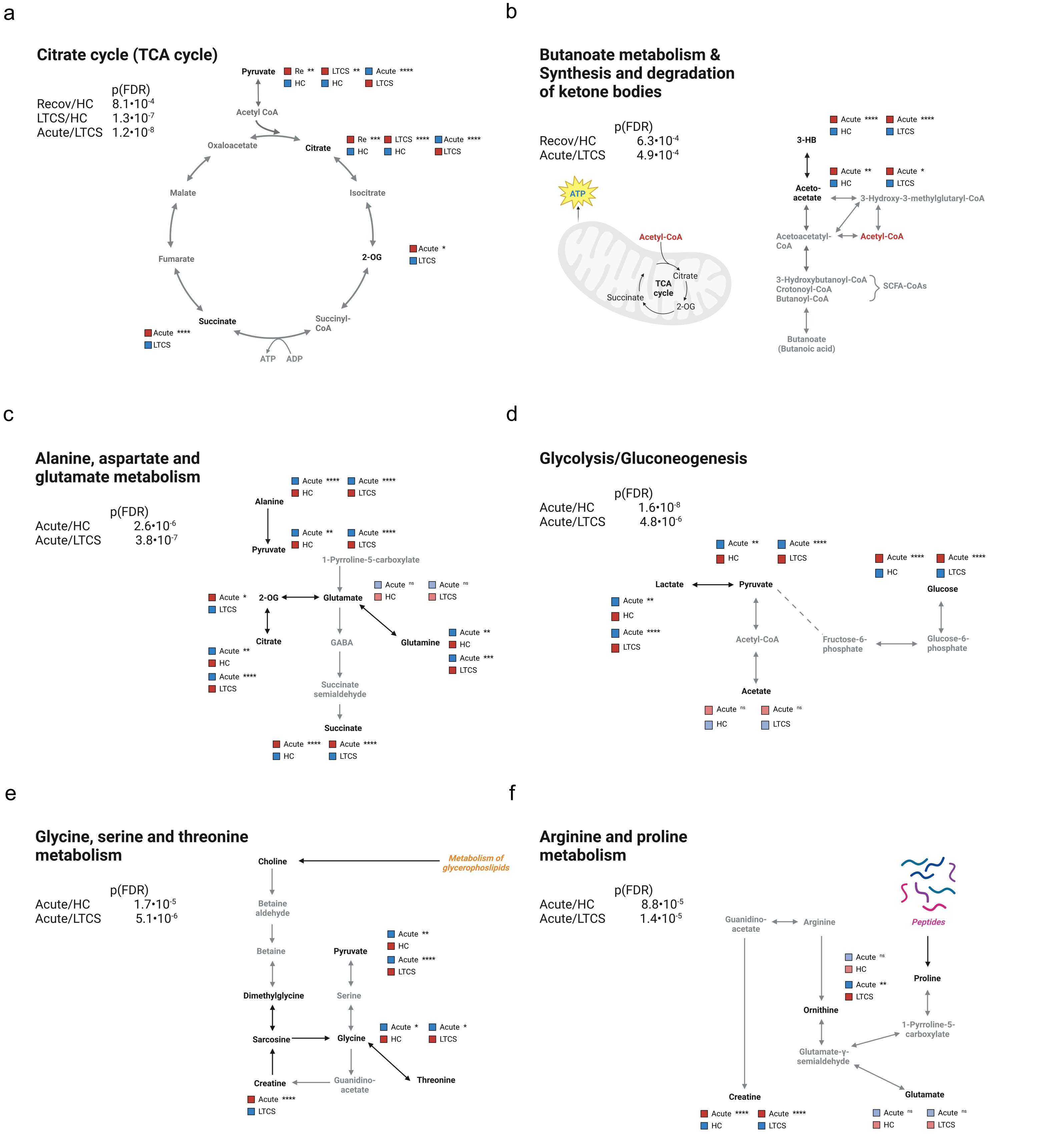

Suppl. Fig. 4

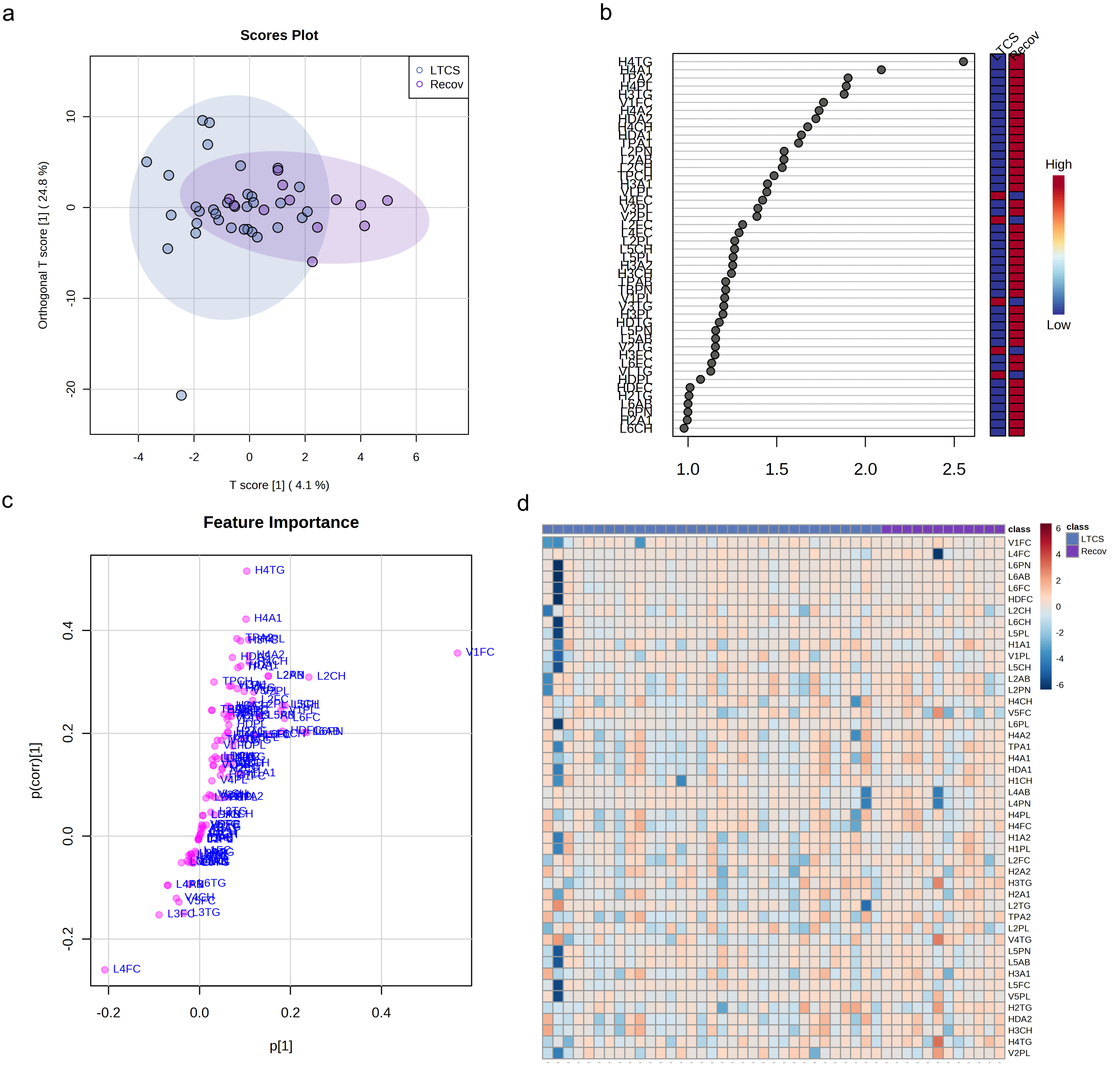

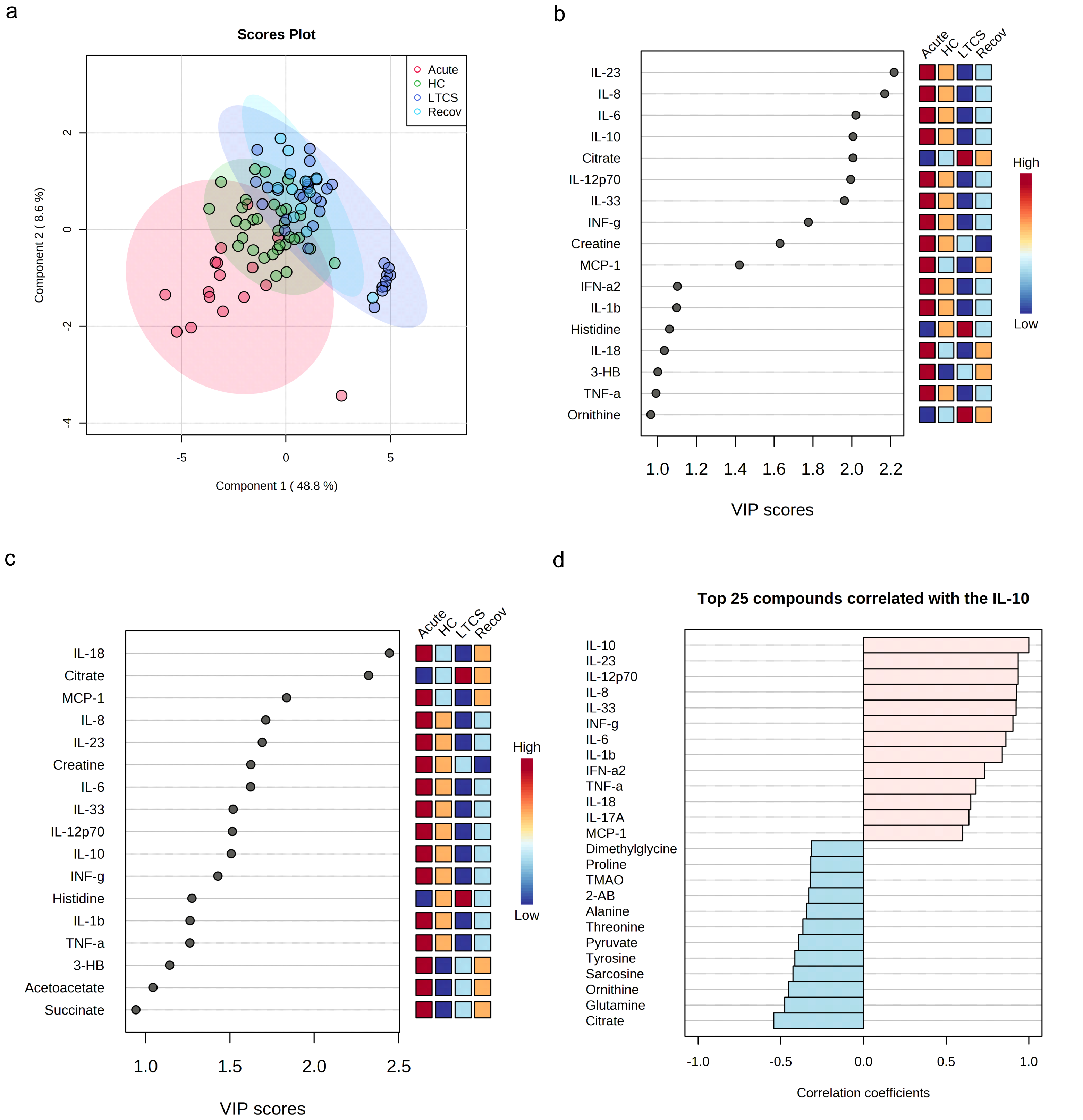
